# Qualification of ELISA and neutralization methodologies to measure SARS-CoV-2 humoral immunity using human clinical samples

**DOI:** 10.1101/2021.07.02.450915

**Authors:** Sasha E. Larsen, Bryan J. Berube, Tiffany Pecor, Evan Cross, Bryan P. Brown, Brittany Williams, Emma Johnson, Pingping Qu, Lauren Carter, Samuel Wrenn, Elizabeth Kepl, Claire Sydeman, Neil P. King, Susan L. Baldwin, Rhea N. Coler

## Abstract

In response to the SARS-CoV-2 pandemic many vaccines have been developed and evaluated in human clinical trials. The humoral immune response magnitude, composition and efficacy of neutralizing SARS-CoV-2 are essential endpoints for these trials. Robust assays that are reproducibly precise, linear, and specific for SARS-CoV-2 antigens would be beneficial for the vaccine pipeline. In this work we describe the methodologies and clinical qualification of three SARS-CoV-2 endpoint assays. We developed and qualified Endpoint titer ELISAs for total IgG, IgG1, IgG3, IgG4, IgM and IgA to evaluate the magnitude of specific responses to the trimeric spike (S) antigen and total IgG specific to the spike receptor binding domain (RBD) of SARS-CoV-2. We also qualified a pseudovirus neutralization assay which evaluates functional antibody titers capable of inhibiting the entry and replication of a lentivirus containing the Spike antigen of SARS-CoV-2. To complete the suite of assays we qualified a plaque reduction neutralization test (PRNT) methodology using the 2019-nCoV/USA-WA1/2020 isolate of SARS-CoV-2 to assess neutralizing titers of antibodies in plasma from normal healthy donors and convalescent COVID-19 individuals.

## 1. Introduction

In December 2019, a series of severe viral pneumonia cases were detected in Wuhan, Hubei Province, China, and later discovered to be induced by a novel coronavirus^1^. Since then, the severe acute respiratory syndrome corona virus 2 (SARS-CoV-2) has spread into a pandemic^2^. As SARS-CoV-2 continues to devastate the world and its economies, the pressure for effective and equitable vaccine distribution remains a global priority^3,4^. Human angiotensin-converting enzyme 2 (ACE2) serves as the host receptor for SARS-CoV-2 surface glycoprotein spike, enabling viral entry into pulmonary alveolar epithelial cells^5-8^. Therefore, many vaccine and therapeutic candidates are designed to induce a robust humoral immune response against SARS-CoV-2 spike protein in an attempt to neutralize the receptor-binding domain (RBD)^9^ and reduce viral entry into host cells upon exposure^10^. This strategy is similar to that employed for SARS-CoV^11^. When measuring the efficacy of therapeutics and vaccines, high-throughput, and clinically qualified reproducible immunological assays provide crucial standardized information regarding the quantity and quality of antibody and cell-mediated responses^12,13^. Specifically, the quantification of antibodies in a spike antigen-based endpoint titer (EPT) enzyme-linked immunosorbent assay (ELISA) in tandem with efficacy measurements via plaque reduction neutralization tests (PRNT) or pseudovirus neutralization, can be leveraged to evaluate responses to SARS-CoV-2 infections and interventions^14-18^.

Indeed, many ELISAs have been developed against SARS-CoV-2^19,20^, including those that characterize seroconversion in response to infection and quantify the magnitude of the humoral response^9^. However, few are clinically qualified using trimeric SARS-CoV-2 spike antigen as our studies report. In order to meet this need, we have developed and qualified a comprehensive suite of high-throughput 384-well format ELISAs for detecting abundance of total human immunoglobulin G (IgG), IgG subclasses (IgG1, IgG3, IgG4), IgM, and IgA recognizing SARS-CoV-2 spike antigen and total IgG specific for RBD. Like other research groups^21,22^ we were unable to identify a sufficient number of samples with robust IgG2 responses to qualify the assay for that subclass. Importantly, we have directly compared our calculated endpoint titer values with WHO international reference standards to make future comparisons between labs more accessible (WHO/BS/2020.2403). In tandem we have qualified clinical SARS-CoV-2 PRNT and pseudovirus neutralization assays evaluating the ability of antibodies in a given sample to reduce viral cell entry as a quality gold-standard surrogate for vaccine-mediated immunogenicity and efficacy^18,23^. Each assay was qualified across precision^24^, linearity^25^ and specificity^13^ endpoints to establish they 1) were accurate and repeatable across users and days, 2) showed a linear response with respect to sample concentrations, and 3) were explicitly detecting responses to SARS-CoV-2. We believe this combination of clinically qualified serology assays will be valuable in comparing responses from various vaccines^26-30^, cutting-edge boost regimens^31,32^, as well as evaluating preclinical vaccine candidates^33,34^ poised to enter clinical trials. Furthermore, these assays will serve to aid the scientific community by allowing the assessment long term vaccine durability^35,36^ and with adaptation the intermediate vaccine efficacy against circulating and emerging viral variants^37^.

## 2. Methods and Materials

### 2.1 Specimens and Controls

Positive controls were obtained through the Seattle Children’s Research Institute’s (SCRI) Center for Global Infectious Disease Research Biorepository. These samples were drawn from Seattle Children’s workforce and adult community members diagnosed with COVID-19 by detection of SARS-CoV-2 nucleic acids in nasopharyngeal specimens, who subsequently enrolled into the Seattle Children’s SARS2 Recovered Cohort with informed consent and Institutional Review Board approval through SCRI. Commercially sourced (Bloodworks Northwest) positive controls were also used from confirmed and recovered COVID-19 patients, termed convalescent plasma. Commercial convalescent plasma was also pooled (CCPP) as a standardized positive control across assays. A sample of pooled plasma from 11 SARS-CoV-2 positive patients from the National Institute for Biological Standards and Controls (NIBSC 20/136), given an arbitrary WHO international standard value of 250 international units (IU) of binding antibody activity per ampule, was also obtained for use in selected assays. Negative plasma originated from historical in-house biorepository of samples isolated from individuals before November 2019, predating the SARS-CoV-2 pandemic and thereby reducing the opportunity for existing cross-reactive antibody responses. Commercially available Normal Human Plasma (Boston Biomedical, Cambridge, MA) was pooled from several donors (NHPP) and used as a standard negative control.

### 2.2 ELISA Methodology

#### Coating

Sterile High Binding 384 well plates (Corning, Corning, NY) were filled with 50 µL per well of ELISA coating buffer containing full-length trimeric SARS-CoV-2 spike protein 11.8 µM or RBD, expressed and purified as described previously^38^, with a final concentration of 1 µg/ml. Coating buffer was made in house using one standard packet of ELISA coating buffer powder (ebioscience, San Diego, CA) mixed with 1 L of distilled water and filtered sterile with a 22 μ m filter to generate 1L of 0.01 M PBS, pH 7.4. Plates were incubated for at least 2 hours at room temperature, or up to 3 days at 4°C.

#### Blocking

Wash Buffer A was made by diluting 1 L of 20X Wash Buffer A solution (Teknova, Hollister, CA) in 19 L of H_2_O from a Barnstead Nanopure Water system (Thermo Scientific, Waltham, MA). After incubation with SARS-CoV-2 Spike protein, plates were washed three times with 100 µL per well of 1X Wash Buffer A. Plates were washed using a BioTek EL406 plate washer (BioTek, Winooski, VT). Blocking buffer was formulated on site by adding 10.0g bovine serum albumin (Sigma, St. Louis, MO) in 1000 mL of phosphate buffered saline (PBS, Lonza, Basel, Switzerland) with 0.05% Tween. Each well received 100 µL of blocking buffer for all plates. Plates were incubated for at least 2 hours, or wrapped in plastic wrap and incubated at 4°C overnight.

#### Sample addition

After blocking, plates were washed three times with 1X Wash Buffer A and subsequently 50 µL of diluent (mixture of 1.0g bovine serum albumin, 500 mL 1X Wash Buffer A and 500 mL PBS) was added to every well of the washed plates. Next, plasma samples were diluted 1:20 in diluent (5 µL of plasma to 95 µL of diluent for total IgG, other Ig sample dilutions listed in **Table 1**) in a 96-well master block before their addition to the 384-well ELISA plate. Using a multichannel pipette, 12.5 µL of the master block sample dilution was manually pipetted into the first column of each sample and subsequently diluted 1:5 from left to right across the plate, discarding 12.5 µL from the final dilution column. Plate layouts varied depending on which aspect of the qualification was being examined. Plates were incubated overnight at 4°C.

**Table 1:**
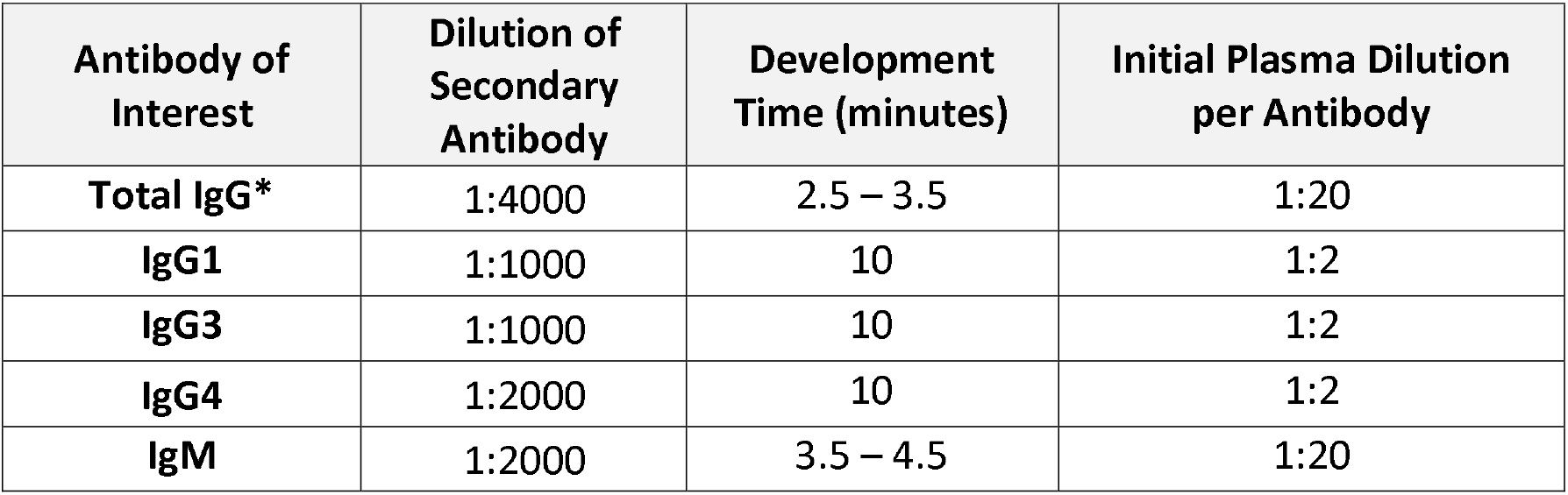
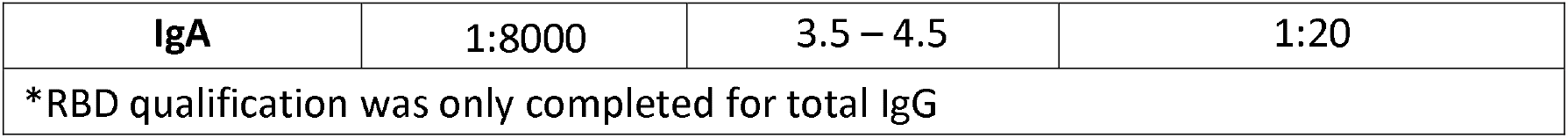
Parameters for each immunoglobulin qualified in ELISA methodology.

| Antibody of Interest | Dilution of Secondary Antibody | Development Time (minutes) | Initial Plasma Dilution per Antibody |
| --- | --- | --- | --- |
| Total IgG* | 1:4000 | 2.5 – 3.5 | 1:20 |
| IgG1 | 1:1000 | 10 | 1:2 |
| IgG3 | 1:1000 | 10 | 1:2 |
| IgG4 | 1:2000 | 10 | 1:2 |
| IgM | 1:2000 | 3.5 – 4.5 | 1:20 |
| **IgA** | 1:8000 | 3.5 – 4.5 | 1:20 |
| *RBD qualification was only completed for total IgG |  |  |  |

#### Development

After incubation, plates were removed from the refrigerator and left to warm to room temperature. Once warmed, plates were washed five times with 1X Wash Buffer A. HRP conjugated rec-Protein G antibodies (Invitrogen, 101223) were diluted (diluent mixture described above) by a factor of 4000, HRP conjugated IgM antibodies (ThermoFisher, A18835) were diluted by a factor of 2000, HRP conjugated IgA antibodies (ThermoFisher, PAI-74395) were diluted by a factor of 8000, HRP conjugated IgG1 antibodies (Invitrogen, MH1715) were diluted by a factor of 1000, HRP conjugated IgG3 antibodies (Invitrogen, 05-3620) were diluted by a factor of 1000, and HRP conjugated IgG4 antibodies (ThermoFisher, MH1742) were diluted by a factor of 2000 **(Table 1)**. Plates were incubated for one hour in the dark at room temperature, then washed five times with 1X Wash Buffer A followed by one wash with PBS. Plates then received 100 µL of Tetramethylbenzidine (TMB) (SeraCare, Milford, MA) per well. After 4-10 minutes; **(Table 1)** depending on secondary antibody, 25 µL of 1N H_2_SO_4_ (Sigma, St. Louis, MO) was added to each well to halt the TMB reaction. Plates were read at a wavelength of 450 nm with a reference filter set at 570 nm using a SpectraMax i3x plate reader (Molecular Devices, San Jose, CA) and SoftMax Pro 6.4.2 analysis software.

#### EPT Calculations

Each diluted sample O.D. value was calculated using 450 nm minus 570 nm values and the average O.D. value for all NHPP dilutions was used to set a minimum cutoff value for each plate. These plate cutoff values were then used to calculate each sample EPT using a 4-parameter logistic model in XL-fit software (model 208) as a Microsoft Excel add-in, as used by our team in previous clinical trial evaluations^39,40^. The EPT value calculated for each duplicate plate is then averaged for a final EPT run value.

#### Variations for competition ELISA

The competition ELISA followed procedures outlined as above except for the following changes. Before adding plasma samples to coated and blocked plates, they were pre-incubated with SARS-CoV-2 Spike protein or RBD at decreasing concentrations. In a 96-well setup plate, soluble SARS-CoV-2 spike or RBD antigens were serially 5-fold diluted. Plasma samples were diluted 1:250 in diluent, and 100 µL was added to the setup plate containing spike protein or RBD. The setup/co-incubation plate was incubated at room temperature for 4 hours with gentle rotation. The plasma/soluble spike or soluble RBD mixtures were added to spike or RBD-coated and blocked plates and the remainder of the ELISA methodology was followed as described above.

### 2.3 Neutralization Methodologies

#### Cell lines and growth conditions

Vero E6 and HEK293T cells were obtained from ATCC and HEK293T cells stably transfected with human Angiotensin-Converting Enzyme 2 (HEK293T-hACE2) were obtained from BEI Resources (NR-52511). All cells were maintained in Dulbecco’s Modified Eagle Media (DMEM) containing 10% fetal bovine serum (FBS), 1% penicillin/streptomycin, and 2 mM L-glutamine (complete DMEM; cDMEM) in a 37°C + 5% CO_2_ incubator. Cells were maintained 25 to 90% confluent and used for the assay up to but not past 14 passages.

#### Pseudovirus production

HIV-1-based SARS-CoV-2 Spike pseudoviruses were prepared as previously described^41^. In brief, HEK293T cells were co-transfected with plasmids in the SARS-CoV-2 Spike-pseudotyped lentiviral particle kit (BEI Resources; NR-52948) using BioT transfection reagent (Bioland Scientific). Plasmids encode for that SARS-CoV-2 Spike protein (Wuhan-Hu-1; Genbank: NC 045512), a lentiviral backbone containing Luc2, and HIV-1 Gag, Pol, Tat1b, and Rev1b. After 48-56 hours, the supernatant was collected and passed through a 0.45 µm filter and aliquots were made and stored at - 80°C until use.

#### Pseudovirus titration and neutralization assay

One day prior to infection, black-walled 96-well plates were coated with 0.01% poly-L-lysine for 5 minutes. Wells were washed twice with sterile water, and HEK293T-hACE2 cells were plated at 2.5×10^4^ cells/well in 50 µL cDMEM. Plates were incubated at 37°C + 5% CO_2_ overnight. To titer the pseudovirus, aliquots were thawed and 2-fold dilutions of pseudovirus in cDMEM were prepared in a 96-well plate. Virus dilutions (100 µL) were added to the HEK293T-hACE2 cells followed immediately by addition of polybrene at a concentration of 5 µg/mL. Cells were incubated at 37°C + 5% CO_2_. After 48 hours, 100 µL of supernatants were removed from the wells, and cells were lysed with 30 µL Bright Glo Luciferase Reagent. Relative luminescence units (RLU) were measured immediately on a Spectramax i3x plate reader, as a surrogate for pseudovirus entry and replication.

For pseudovirus neutralization assays, plasma samples were diluted in cDMEM in a 10-point 3-fold dilution series in a 96-well setup plate. SARS-CoV-2 Spike pseudovirus was diluted to 1×10^5^ RLU/mL and added to wells of the setup plate in a 1:1 ratio. The setup plate was incubated at 37°C + 5% CO_2_ for 1 hour prior to addition to HEK293T-hACEs cell as above. Percent inhibition was calculated by dividing RLU values in each well to the average RLU value of virus only wells. Pseudovirus inhibition curves and the concentration of plasma required to inhibit pseudovirus entry by 50% (IC_50_) was determined by plotting the data and fitting a curve using the neutcurve Python package (https://jbloomlab.github.io/neutcurve/).

#### SARS-CoV-2 virus production

SARS-CoV-2 isolate 2019-nCoV/USA-WA1/2020 was obtained from BEI Resources and housed under standard BSL-3 laboratory conditions. SARS-CoV-2 virus was propagated and titered by plaque assay in Vero E6 cells. The original stock virus was added to cells at an MOI of 0.1 and incubated at 37°C + 5% CO_2_ for 72 hours. Supernatants were harvested, spun at 1,200 rpm for 20 minutes, and the supernatant was aliquoted and frozen at -80°C to create a Passage 1 (P1) virus stock. P1 virus was titered and propagated a second time as above to create Passage 2 (P2) virus. Aliquots of the P2 virus were diluted to 4.5 × 10^3^ plaque-forming units (PFU)/mL to create a PRNT virus stock.

#### PRNT assay

One day prior to infection, Vero E6 cells were plated at 4 × 10^5^ cells/well in 2 mL cDMEM in a 6-well plate. The following day, heat-inactivated serum/plasma samples (incubated at 56 °C for 1 hour) were diluted in dilution media (DMEM + 1% FBS) in a 6-point 2-fold dilution series in a 96-well setup plate to a total volume of 115 µL per well. SARS-CoV-2 PRNT stocks were thawed and diluted 5-fold in dilution media and 115 µL added to the setup plate (9.0 × 10^2^ PFU/mL). Virus and plasma were co-incubated for 1 hour at 37°C + 5% CO_2_. Media was carefully removed from the 6-well plates and 200 µL of the virus/plasma mixture was added to the cells. The plates were incubated at 37°C + 5% CO_2_ with rocking every 15 minutes. After 1 hour, 2.0 mL overlay media (dilution media + 0.2% agarose, kept at a minimum of 45°C) was added to the wells and plates were subsequently incubated at 37°C + 5% CO_2_ for 72 hours. After 72 hours, cells were fixed by the addition of 2.0 mL of 10% formaldehyde and incubated for 30 minutes. Fixative and overlay were removed, wells were stained for 20 minutes with 1.0 mL of 0.3% Crystal Violet (BD) and washed once with 1.0 mL PBS per well. Plates were inverted and plaques indicating cell lysis from viral infection were counted for each well and recorded. The lowest dilution of plasma to reduce PFU by 80% compared to virus only wells was deemed the PRNT_80_. The calculation for the threshold is as follows

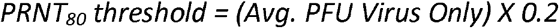

### 2.4 Qualification

**Precision** is the accuracy and reproducibility of an assay across different variables, including between users and days of execution. Given sample endpoint and neutralization titers are unknown, they are not compared to a known standard EPT value but rather evaluated for a consistent performance of the assay to return a similar value for specific control samples across conditions. <u>Intra-assay</u> variability measures the variation of the assay under the same conditions, while <u>inter-assay</u> variability measures variation between different operators and days. Intra-assay variability was measured across three runs performed by a single technician on the same day with like samples. Inter-day variability was measured across three runs performed by a single technician across three separate days with like samples. Finally, for inter-operator variability each of three operators generated their own independent plates and reagents and performed the assay on the same day with like samples. Plate layouts and samples used remained consistent across each measurement of variability. This evaluates the reproducibility of the assay and demonstrates that the procedure generates consistent results.

### ELISA Precision Endpoint

Each sample must return a coefficient of variation (CV) no greater than 20% for any variable (operator, day, run) evaluated. Coefficient of variation is calculated as

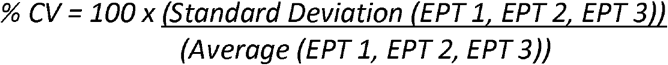

where EPT 1-3 are the calculated endpoint values for an individual sample across the three runs (operator, day) being evaluated. For example, CCPP sample EPT values across each of the three operator runs must not exceed 20% CV. This value is partially based on European Medicines Agency (EMA) guidelines that state precision and repeatability must be below this 20% CV threshold.

### Pseudovirus Neutralization Precision Endpoint

The average from duplicate samples on any given plate must be within 2-fold (+/-) of the geometric mean within the respective category (operator, run, reproducibility). For example, the average IC_50_ value from duplicate samples for CCPP on each plate within an intra-operator variability test must be within 2-fold of the geometric mean IC_50_ value of all CCPP samples tested within the intra-operator variability test.

### PRNT Precision Endpoint

Each sample on any given plate must be within 2-fold (+/-) of the geometric mean within the respective category. For example, each PRNT_80_ value for CCPP within an intra-operator variability test must be within 2-fold of the geometric mean PRNT_80_ value of all CCPP samples tested within the intra-operatory variability test.

**Linearity** helps evaluate how well an assay is able to discern variable concentrations of an analyte and if the detected quantity well represents a value across a dilution series. These samples do not have a reference ‘true value’ so instead we leveraged known positive samples across a standardized dilution curve to determine if the detected response was proportional to the dilution. For ELISA assays, curves were created using O.D. values for positive samples from precision plates and analyzed by linear regression^25^ with calculated R^2^ values for individual samples as well as an aggregate analysis of an adjusted multiple R^2^ following established calculation methods^42^. Only samples with three or more data points were included. Similarly, for PRNT and pseudovirus neutralization assays precision plate dilutions were used from positive samples analyzed by linear regression^25^ with calculated R^2^ values.

### Linearity Endpoint

A graphical analysis of the line of best fit (by least squares criterion) coupled with an R^2^ value of 0.90 or greater by linear regression analysis, demonstrates the acceptable linearity of the assay.

**Specificity** is determined in two distinct ways to ensure SARS-CoV-2 antigen-specific responses are being captured, including: a robust screening of presumed negatives, and competition assays with soluble antigen pre-competing for antibody binding. This helps promote that responses are not due to cross-reactive humoral responses that are present in unexposed blood donors. A set of 92 (used for each SARS-CoV-2 spike antibody ELISA) or 82 (used for total IgG RBD) negative samples from a pre-COVID-19 biorepository and four positive SARS-CoV-2 convalescent samples (2 positives for IgG4) were used in determining specificity of our ELISA methodology. For total IgG, IgA and IgM ELISAs, samples were diluted 5-fold starting at 1:100 for a 4-point dilution. For IgG1, IgG3, and IgG4 ELISAs, samples were diluted 5-fold starting at 1:10 for a 4-point dilution. A single plate was run by one technician. A competitive ELISA was also performed to further demonstrate specificity of the immune response to SARS-CoV-2. In this specificity assay, we are expecting serum or plasma antibodies to bind and be sequestered from the coated plate by higher concentrations of soluble spike or RBD in solution. A positive sample was run in duplicate on the same plate by a single operator for this analysis.

We evaluated the specificity of serum/plasma samples to neutralize SARS-CoV-2 via a robust negative sample size. We used 20 negative plasma samples from a historical (pre-COVID) biorepository and 4 positive control samples from patients who became infected and recovered from COVID-19 as a characterization of the pseudovirus neutralization and PRNT assay specificity. For pseudovirus neutralization, each sample was assessed at a 1:20 dilution and was run in duplicate by a single operator on the same day. In addition, we tested plasma from a convalescent patient, as well as a pool of plasma from a set of convalescent patients, to evaluate specificity against a non-specific pseudovirus harboring the VSV G cell entry protein instead of the SARS-CoV-2 spike protein. For PRNT specificity, 20 negative samples from a pre-COVID-19 biorepository and four positive samples were used. Each sample was assessed at a 1:40 dilution for and run in duplicate by a single operator.

### ELISA Specificity Endpoints

For ELISAs ≥ 90% of negative sample O.D.s must be lower than the collective average of all negative samples + 3 SD (plate limit of detection [LOD]), while all 4 positives must exceed this value. For the competition ELISA the results must demonstrate that as the amount of exogenous SARS-CoV-2 Spike antigen or RBD decreases, there will be an increase in positive signal measured by O.D. 450nm-570nm. Graphical analysis of O.D. values versus dilution values and a linear relationship with an R^2^ value > 0.90 demonstrate an acceptable competition specificity qualification of the ELISA. Samples are recorded as pass-fail for the negative-screening specificity criteria.

### Neutralization Specificity Endpoints

For pseudovirus neutralization specificity, all 20 negative samples must not achieve an IC_50_ value at the 1:20 dilution of plasma, whereas all 4 positive samples must meet this threshold. Samples are recorded as pass-fail for these criteria. For the VSV G pseudovirus specificity test, all COVID-positive samples must not achieve an IC_50_ value at the 1:20 dilution of plasma. For PRNT, the observed PFU from all 20 negative samples must not achieve a PRNT_80_ at the 1:40 dilution of plasma, whereas all 4 positive samples must meet this threshold.

## 3 Results and Discussion

### 3.1 ELISA

#### 3.1.1 Precision

Assay repeatability, or the degree to which an assay will produce the same result when identical test samples are evaluated under the same operating conditions, was assessed in terms of run to run variation. Samples from 14 subjects and 2 pooled controls (CCPP and NHPP) were run by a single operator in triplicate across three separate days (interday), by a single operator in triplicate on the same day (intraday), and by three individual operators on the same day (interoperator). Representative data for Total IgG SARS-CoV-2 spike antigen-specific responses for each precision test is shown in **Figure 1** including two individual convalescent samples (M and N) and pooled controls (CCPP, NHPP). **Figure 2** shows representative samples assess for SARS-CoV-2 RBD responses across the different variables. The CV for each sample (16 total) across the precision variables evaluated are reported in **Tables 2, 3 and 4**, and cumulative EPT graphs containing all data points are in **Supplemental Figure 1**. Importantly, across the robust set of samples tested, all were below the 20% CV endpoint established for this assay, demonstrating the assay is reproducible and precise.

**Figure 1:**
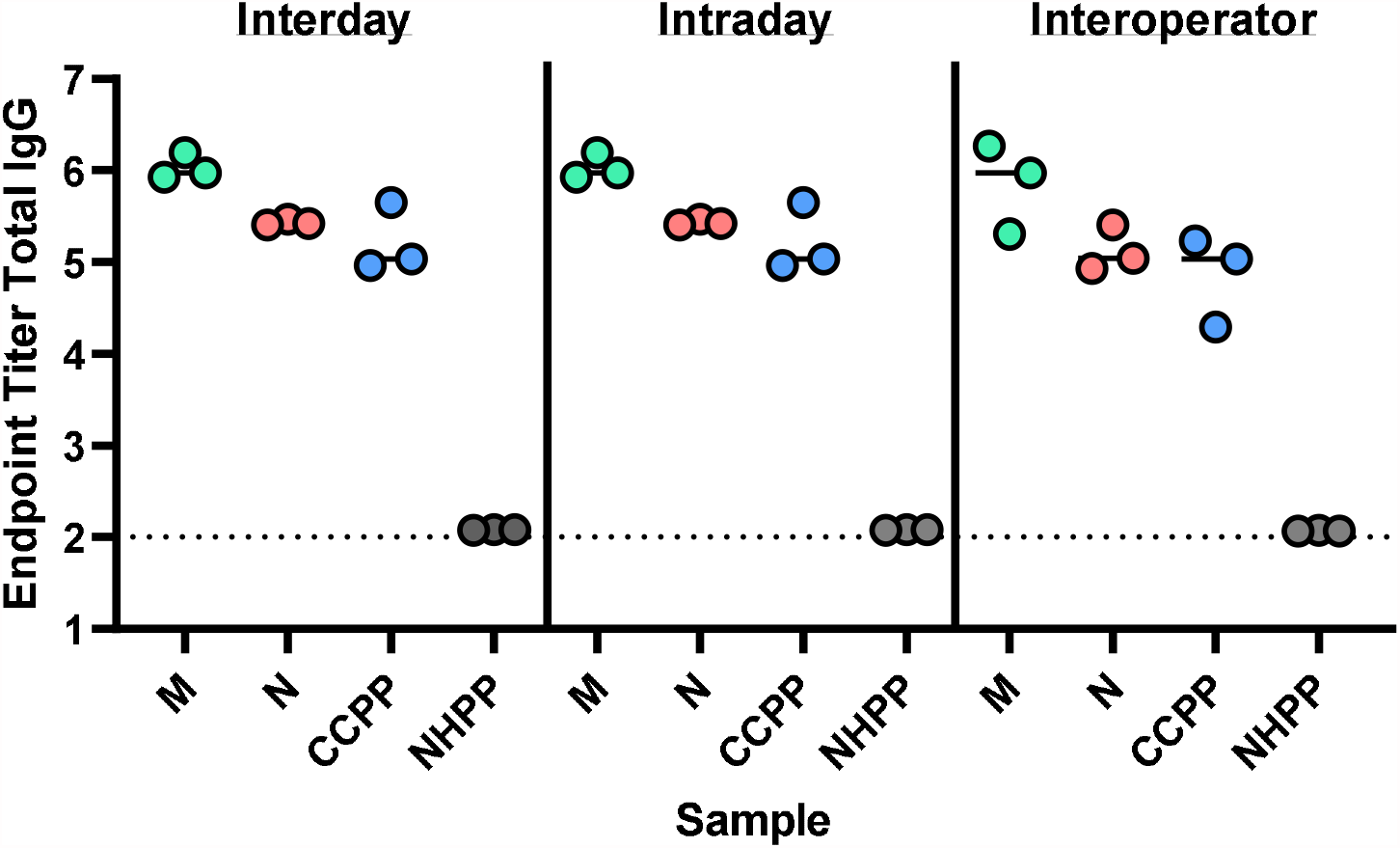
SARS-CoV-2 Spike Total IgG Endpoint Titers. evaluated for two individual samples (M, N) as well as pooled positive (CCPP) and negative (NHPP) control samples.

**Figure 2:**
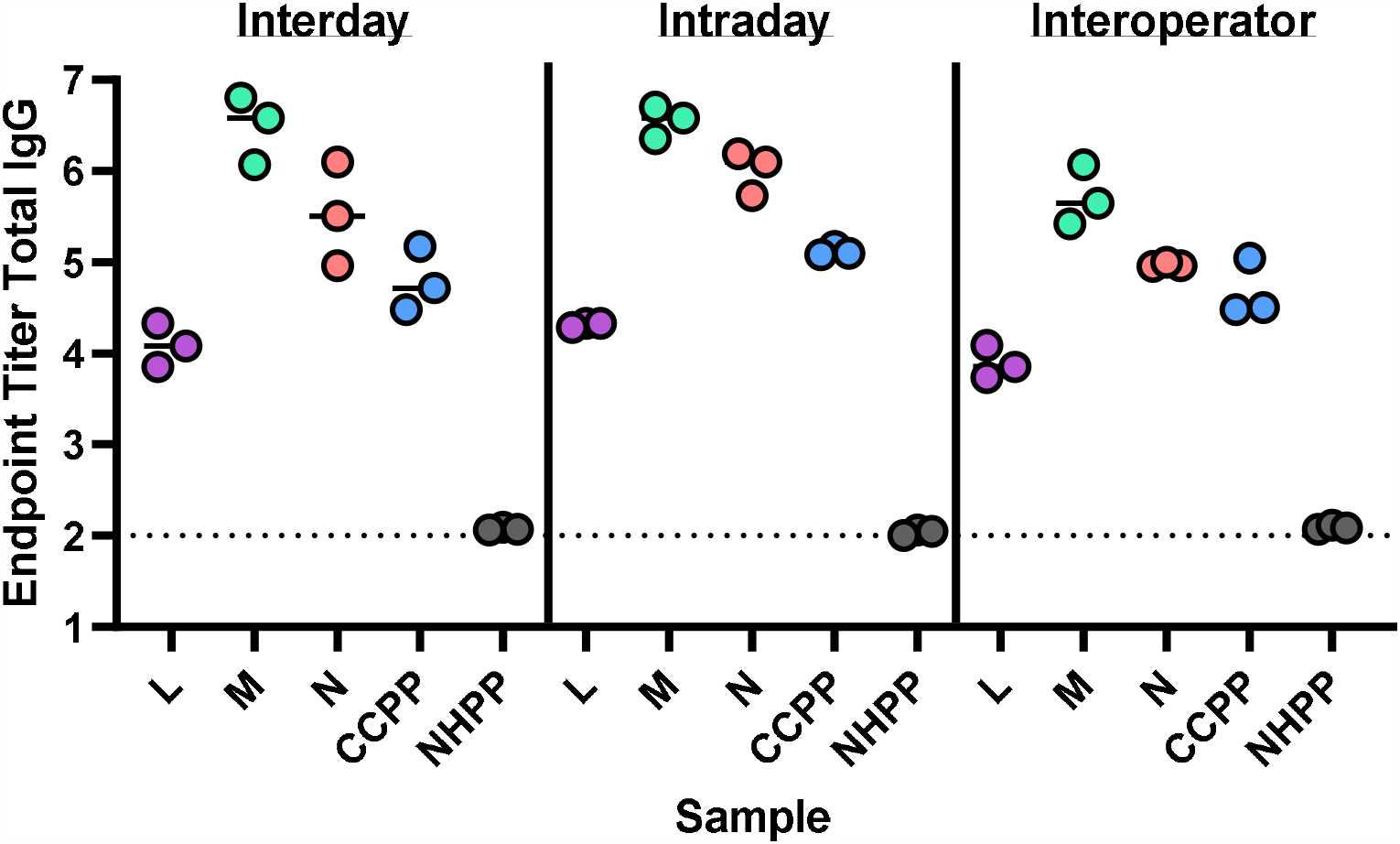
SARS-CoV-2 RBD Total IgG Endpoint Titers. evaluated for three individual samples (L, M, N) as well as pooled positive (CCPP) and negative (NHPP) control samples.

**Table 2:**
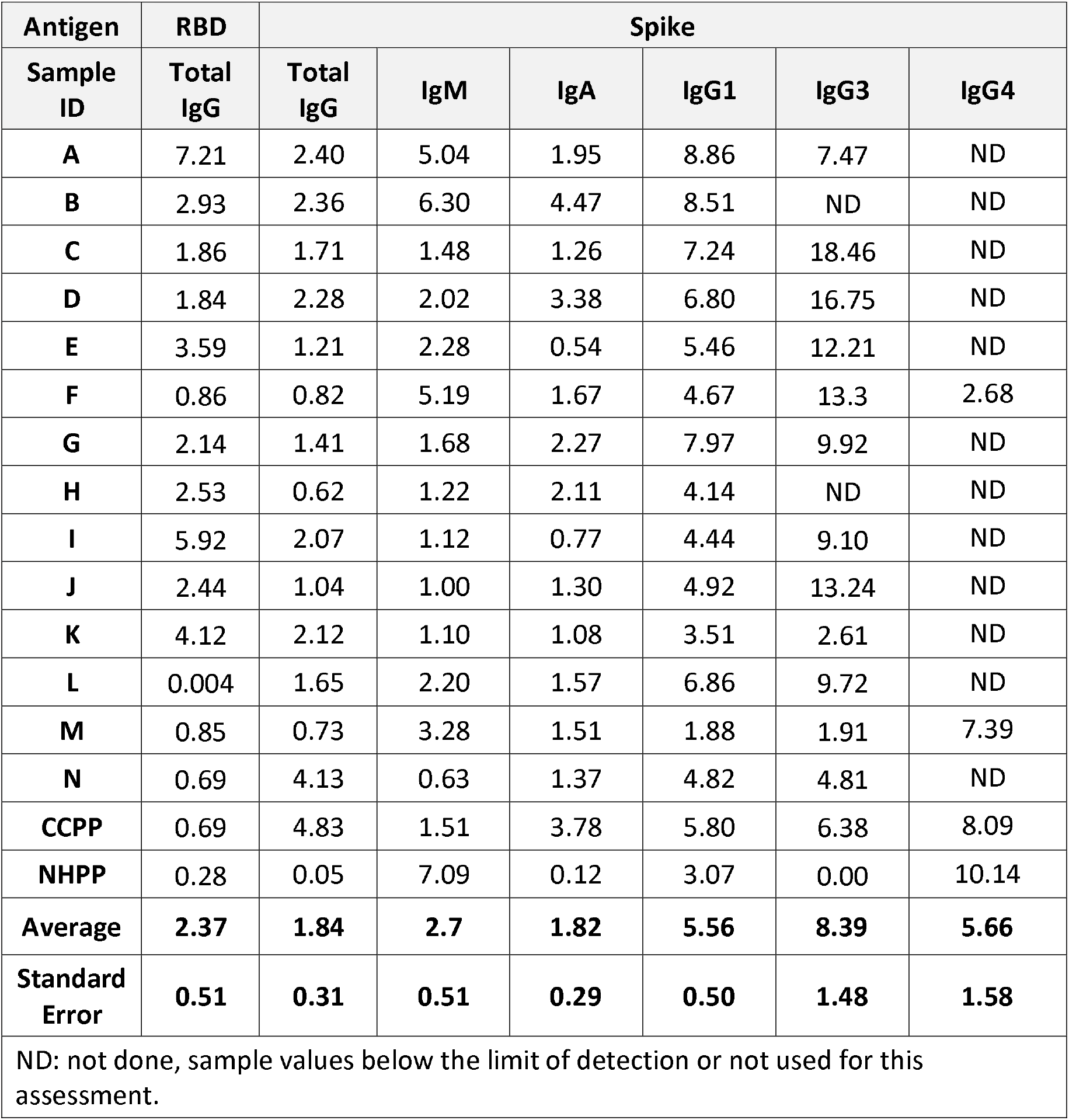
ELISA Intraday Precision analysis using coefficient of variation (%)

| Antigen | RBD | Spike |  |  |  |  |  |
| --- | --- | --- | --- | --- | --- | --- | --- |
| Sample ID | Total IgG | Total IgG | IgM | IgA | IgG1 | IgG3 | IgG4 |
| A | 7.21 | 2.40 | 5.04 | 1.95 | 8.86 | 7.47 | ND |
| B | 2.93 | 2.36 | 6.30 | 4.47 | 8.51 | ND | ND |
| C | 1.86 | 1.71 | 1.48 | 1.26 | 7.24 | 18.46 | ND |
| D | 1.84 | 2.28 | 2.02 | 3.38 | 6.80 | 16.75 | ND |
| E | 3.59 | 1.21 | 2.28 | 0.54 | 5.46 | 12.21 | ND |
| F | 0.86 | 0.82 | 5.19 | 1.67 | 4.67 | 13.3 | 2.68 |
| G | 2.14 | 1.41 | 1.68 | 2.27 | 7.97 | 9.92 | ND |
| H | 2.53 | 0.62 | 1.22 | 2.11 | 4.14 | ND | ND |
| I | 5.92 | 2.07 | 1.12 | 0.77 | 4.44 | 9.10 | ND |
| J | 2.44 | 1.04 | 1.00 | 1.30 | 4.92 | 13.24 | ND |
| K | 4.12 | 2.12 | 1.10 | 1.08 | 3.51 | 2.61 | ND |
| L | 0.004 | 1.65 | 2.20 | 1.57 | 6.86 | 9.72 | ND |
| M | 0.85 | 0.73 | 3.28 | 1.51 | 1.88 | 1.91 | 7.39 |
| N | 0.69 | 4.13 | 0.63 | 1.37 | 4.82 | 4.81 | ND |
| CCPP | 0.69 | 4.83 | 1.51 | 3.78 | 5.80 | 6.38 | 8.09 |
| NHPP | 0.28 | 0.05 | 7.09 | 0.12 | 3.07 | 0.00 | 10.14 |
| Average | 2.37 | 1.84 | 2.7 | 1.82 | 5.56 | 8.39 | 5.66 |
| Standard Error | 0.51 | 0.31 | 0.51 | 0.29 | 0.50 | 1.48 | 1.58 |
| ND: not done, sample values below the limit of detection or not used for this assessment. |  |  |  |  |  |  |  |

**Table 3:** ELISA Interday Precision analysis using coefficient of variation (%)

| Antigen | RBD | Spike |  |  |  |  |  |
| --- | --- | --- | --- | --- | --- | --- | --- |
| Sample ID | Total IgG | Total IgG | IgM | IgA | IgG1 | IgG3 | IgG4 |
| A | 5.94 | 2.24 | 8.71 | 3.24 | 10.69 | 12.94 | ND |
| B | 6.70 | 2.64 | 3.24 | 2.28 | 13.26 | 4.56 | ND |
| C | 7.50 | 3.71 | 12.92 | 3.80 | 3.29 | 9.90 | ND |
| D | 3.39 | 0.82 | 7.73 | 2.23 | 6.89 | 17.39 | ND |
| E | 5.42 | 0.99 | 10.36 | 1.95 | 4.06 | 8.76 | ND |
| F | 9.50 | 3.03 | 13.40 | 1.56 | 5.11 | 17.68 | 2.12 |
| G | 3.45 | 2.82 | 9.84 | 0.10 | 3.73 | 2.43 | ND |
| H | 3.67 | 1.79 | 3.70 | 0.97 | ND | ND | ND |
| I | 1.63 | 0.91 | 7.98 | 2.19 | 10.65 | 13.21 | ND |
| J | 5.28 | 0.29 | 10.29 | 2.28 | 6.21 | 17.39 | ND |
| K | 6.14 | 4.31 | 9.47 | 3.06 | 2.80 | 4.88 | ND |
| L | 4.72 | 1.41 | 5.06 | 1.53 | 8.76 | 6.69 | ND |
| M | 4.73 | 1.96 | 12.93 | 4.30 | 7.12 | 10.73 | 5.10 |
| N | 8.40 | 0.50 | 6.28 | 2.43 | 8.28 | 13.12 | ND |
| CCPP | 6.00 | 5.86 | 8.81 | 6.02 | 8.54 | 9.80 | ND |
| NHPP | 0.69 | 0.065 | 6.07 | 1.45 | 2.71 | 3.57 | 7.44 |
| Average | 5.20 | 2.08 | 8.55 | 2.46 | 6.81 | 10.20 | 18.98 |
| Standard Error | 0.58 | 0.40 | 0.78 | 0.35 | 0.83 | 1.32 | 1.54 |
| ND: not done, sample values below the limit of detection or not used for this assessment. |  |  |  |  |  |  |  |

**Table 4:**
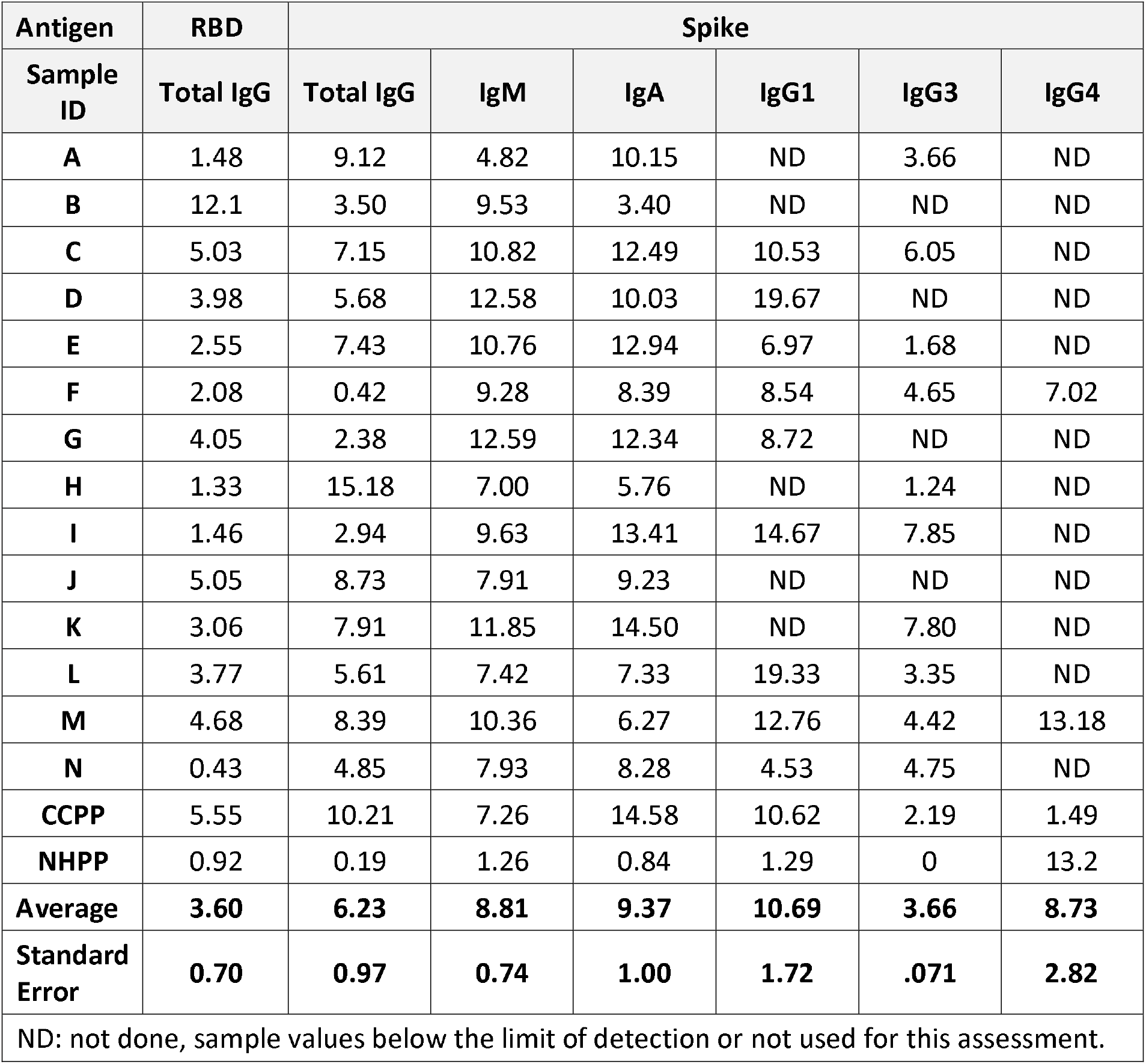
ELISA Interoperator Precision analysis using coefficient of variation (%)

| Antigen | RBD | Spike |  |  |  |  |  |
| --- | --- | --- | --- | --- | --- | --- | --- |
| Sample ID | Total IgG | Total IgG | IgM | IgA | IgG1 | IgG3 | IgG4 |
| A | 1.48 | 9.12 | 4.82 | 10.15 | ND | 3.66 | ND |
| B | 12.1 | 3.50 | 9.53 | 3.40 | ND | ND | ND |
| C | 5.03 | 7.15 | 10.82 | 12.49 | 10.53 | 6.05 | ND |
| D | 3.98 | 5.68 | 12.58 | 10.03 | 19.67 | ND | ND |
| E | 2.55 | 7.43 | 10.76 | 12.94 | 6.97 | 1.68 | ND |
| F | 2.08 | 0.42 | 9.28 | 8.39 | 8.54 | 4.65 | 7.02 |
| G | 4.05 | 2.38 | 12.59 | 12.34 | 8.72 | ND | ND |
| H | 1.33 | 15.18 | 7.00 | 5.76 | ND | 1.24 | ND |
| I | 1.46 | 2.94 | 9.63 | 13.41 | 14.67 | 7.85 | ND |
| J | 5.05 | 8.73 | 7.91 | 9.23 | ND | ND | ND |
| K | 3.06 | 7.91 | 11.85 | 14.50 | ND | 7.80 | ND |
| L | 3.77 | 5.61 | 7.42 | 7.33 | 19.33 | 3.35 | ND |
| M | 4.68 | 8.39 | 10.36 | 6.27 | 12.76 | 4.42 | 13.18 |
| N | 0.43 | 4.85 | 7.93 | 8.28 | 4.53 | 4.75 | ND |
| CCPP | 5.55 | 10.21 | 7.26 | 14.58 | 10.62 | 2.19 | 1.49 |
| NHPP | 0.92 | 0.19 | 1.26 | 0.84 | 1.29 | 0 | 13.2 |
| Average | 3.60 | 6.23 | 8.81 | 9.37 | 10.69 | 3.66 | 8.73 |
| Standard Error | 0.70 | 0.97 | 0.74 | 1.00 | 1.72 | .071 | 2.82 |
| ND: not done, sample values below the limit of detection or not used for this assessment. |  |  |  |  |  |  |  |

#### Intraday variability

One operator completed three runs, totaling three sets of duplicate plates on the same day using like samples (Sample ID A-N, CCPP and NHPP). Endpoint titers were calculated as described above to evaluate the magnitude of total IgG, IgM, IgA, IgG1, IgG3 and IgG4 present in each sample **(Supplemental Figure 2)** and the CV was evaluated across the three runs executed **(Table 2)**. Importantly, the % CV for all samples tested and the average across samples was well below our endpoint of < 20% variability **(Table 2)**.

#### Interday variability

One operator performed three runs with like samples over three different days (Sample ID A-N, CCPP and NHPP). Endpoint titers were calculated as described (**Supplemental Figure 3**) and the CV was evaluated across the three runs executed (**Table 3**). Importantly, all samples and the average across samples %CV was well below our endpoint of < 20% variability (**Table 3**)

#### Interoperator variability

Three different technicians ran two duplicate plates using like samples on the same day. Similarly, endpoint titers were calculated (**Supplemental Figure 4**) and CV evaluated for each antibody ELISA across the three runs executed (**Table 4**). These data demonstrate that the EPT ELISA methodology yields results within an acceptable range of variance, all below 20%, meeting the EMA guidelines.

#### 3.1.2 Linearity

We used a linear regression analysis of O.D. compared to dilution to calculate the R^2^ for a positive samples across each antibody ELISA evaluated for spike and RBD (Total IgG only) responses. Importantly, only sample containing three data points in the linear range were used to calculate R^2^, and those values in plateau ranges were excluded **(Figure 5)**. The adjusted multiple R^2^ values calculated for spike antigen-specific responses were as follows: total IgG 0.930, IgM 0.862, IgA 0.927, IgG1 0.863, and IgG3 0.882 **(Figure 5)**. Given the low responder rate for IgG4 linearity requirements for sample inclusion were not met. The adjusted multiple R^2^ value calculated for total IgG RBD-specific responses was 0.903. Adjusted multiple R^2^ values for spike specific IgA, total IgG and RBD specific total IgG responses exceeded the 0.90 linearity qualification endpoint, demonstrating acceptable linearity for this methodology. While spike specific IgM, IgG3 and IgG1 adjusted multiple R2 values did not meet 0.90, they did approach this threshold. Importantly, the positive plate control CCPP exceeded an R^2^ value of 0.90 for each antibody class, except IgG4, suggesting these assays are linear.

**Figure 5:**
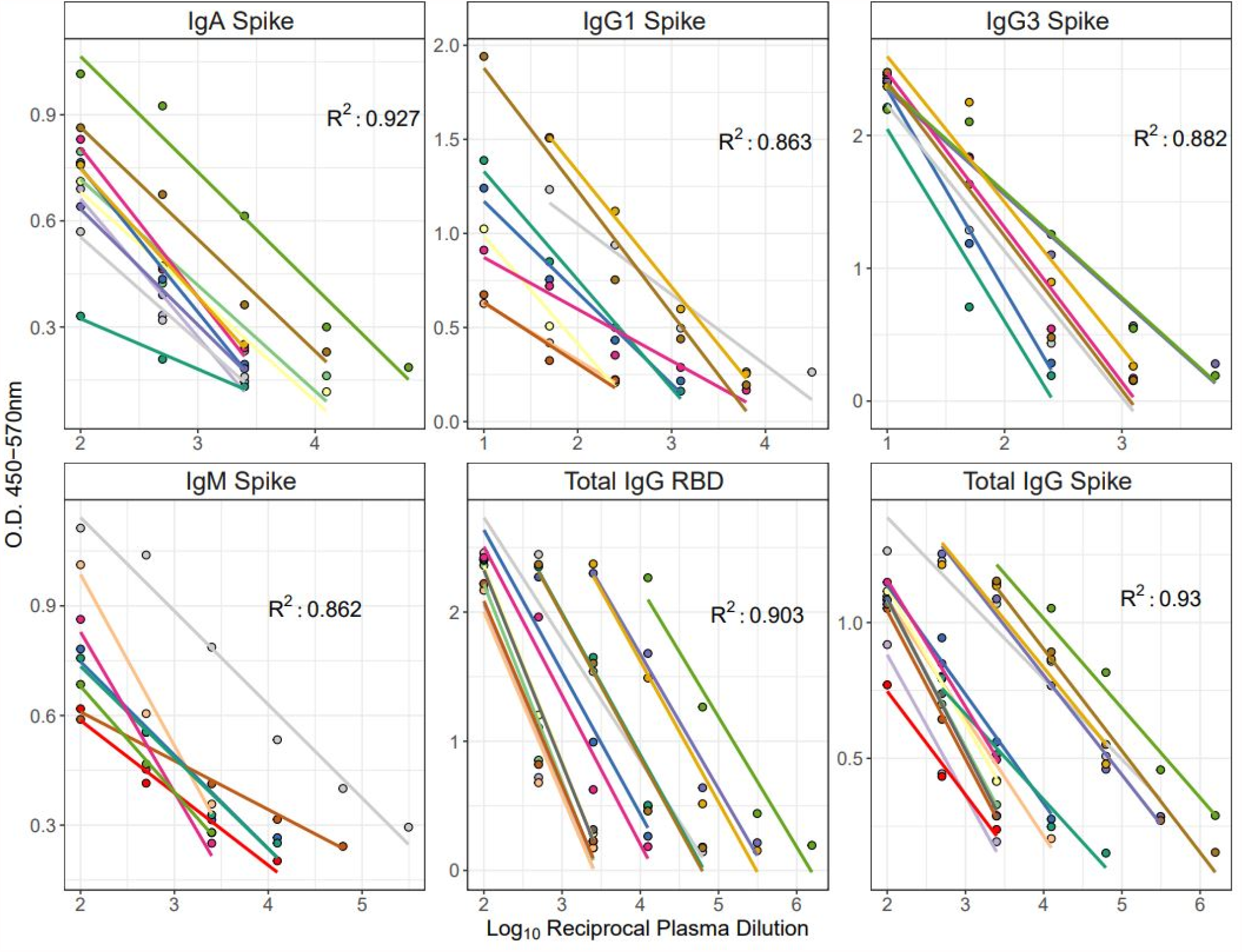
ELISA Linearity for SARS-CoV-2 Spike Responses. Linear regression comparing O.D. 450-570nm values to dilution of plasma for each antibody evaluated is shown. Linear model lines for each sample included for IgA, IgG1, IgG3, IgM and total IgG for both spike and RBD. Each line of best fit represents a unique sample. The adjusted multiple R^2^ value for each antibody class is shown.

#### 3.1.3 Specificity

We evaluated the effectiveness of our assay to specifically distinguish antibodies in samples that recognize SARS-CoV-2 spike or RBD. Each antibody was screened against 92 total presumed negative (pre-pandemic) samples and 4 convalescent samples (or 2 in the case of IgG4) for spike antigen **(Table 5)**, while 82 negative samples were leveraged for RBD ELISA analysis **(Table 6)**. Each antibody evaluated met our endpoint of having ≥ 90% of negative samples screened return true negative values (below 3 SD of the plate LOD, while all SARS-CoV-2 convalescent samples evaluated returned 100% positive values **(Table 5 and Table 6)**, establishing the specificity of our methodology.

**Table 5:**
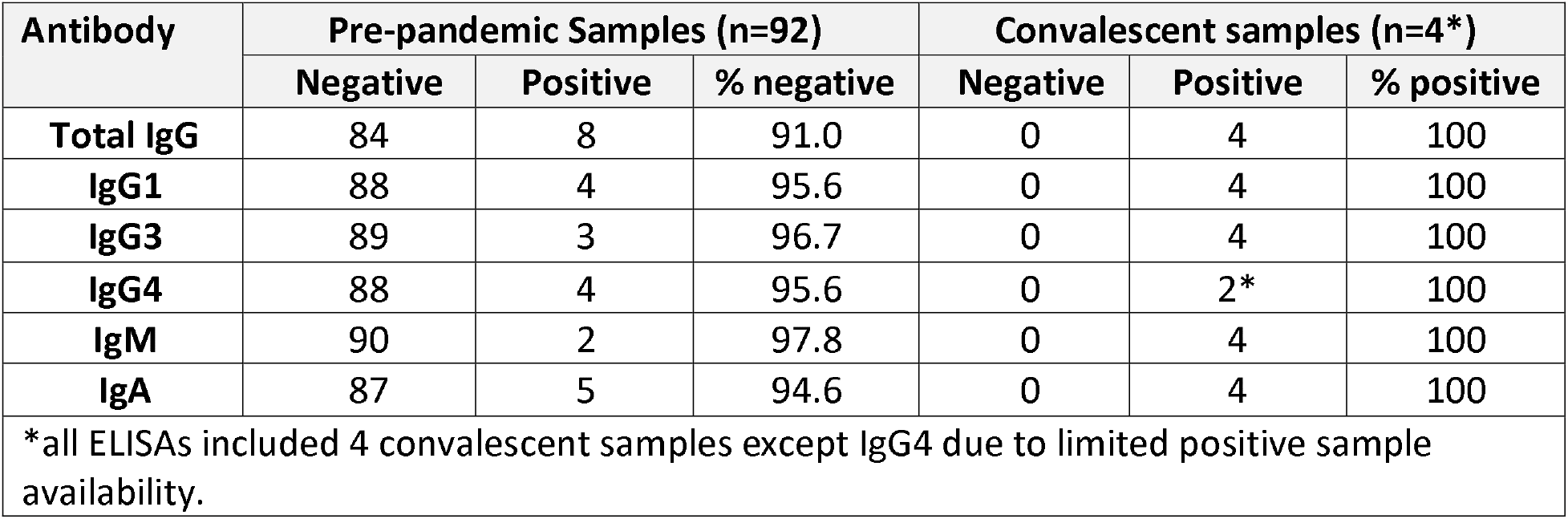
Robust Negative Sample Screen for SARS-CoV-2 Spike ELISA EPT Specificity.

| Antibody | Pre-pandemic Samples (n=92) |  |  | Convalescent samples (n=4*) |  |  |
| --- | --- | --- | --- | --- | --- | --- |
|  | Negative | Positive | % negative | Negative | Positive | % positive |
| Total IgG | 84 | 8 | 91.0 | 0 | 4 | 100 |
| IgG1 | 88 | 4 | 95.6 | 0 | 4 | 100 |
| IgG3 | 89 | 3 | 96.7 | 0 | 4 | 100 |
| IgG4 | 88 | 4 | 95.6 | 0 | 2* | 100 |
| IgM | 90 | 2 | 97.8 | 0 | 4 | 100 |
| IgA | 87 | 5 | 94.6 | 0 | 4 | 100 |
| *all ELISAs included 4 convalescent samples except IgG4 due to limited positive sample availability. |  |  |  |  |  |  |

**Table 6:**
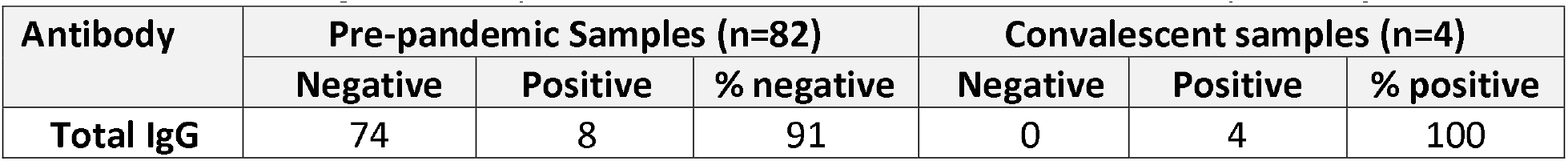
Robust Negative Sample Screen for SARS-CoV-2 RBD ELISA EPT Specificity.

| Antibody | Pre-pandemic Samples (n=82) |  |  | Convalescent samples (n=4) |  |  |
| --- | --- | --- | --- | --- | --- | --- |
|  | Negative | Positive | % negative | Negative | Positive | % positive |
| Total IgG | 74 | 8 | 91 | 0 | 4 | 100 |

#### 3.1.4 Correlation to WHO Reference Standard

We ran a WHO reference positive standard (NIBSC 20/136) with an assigned antibody binding activity of 250 IU per ampule alongside a set of samples used for qualification to equate IU antibody binding activity to EPT. We observed a Total IgG endpoint titer of 4.73 and 4.47 against SARS-CoV-2 trimeric spike and RBD, respectively for the WHO sample **(Figure 6)**. Not surprisingly we saw a consistently lower EPT against RBD than Spike for the same plasma samples, including pooled controls (CCPP and NHPP) as well as the WHO sample **(Figure 6)**.

**Figure 6:**
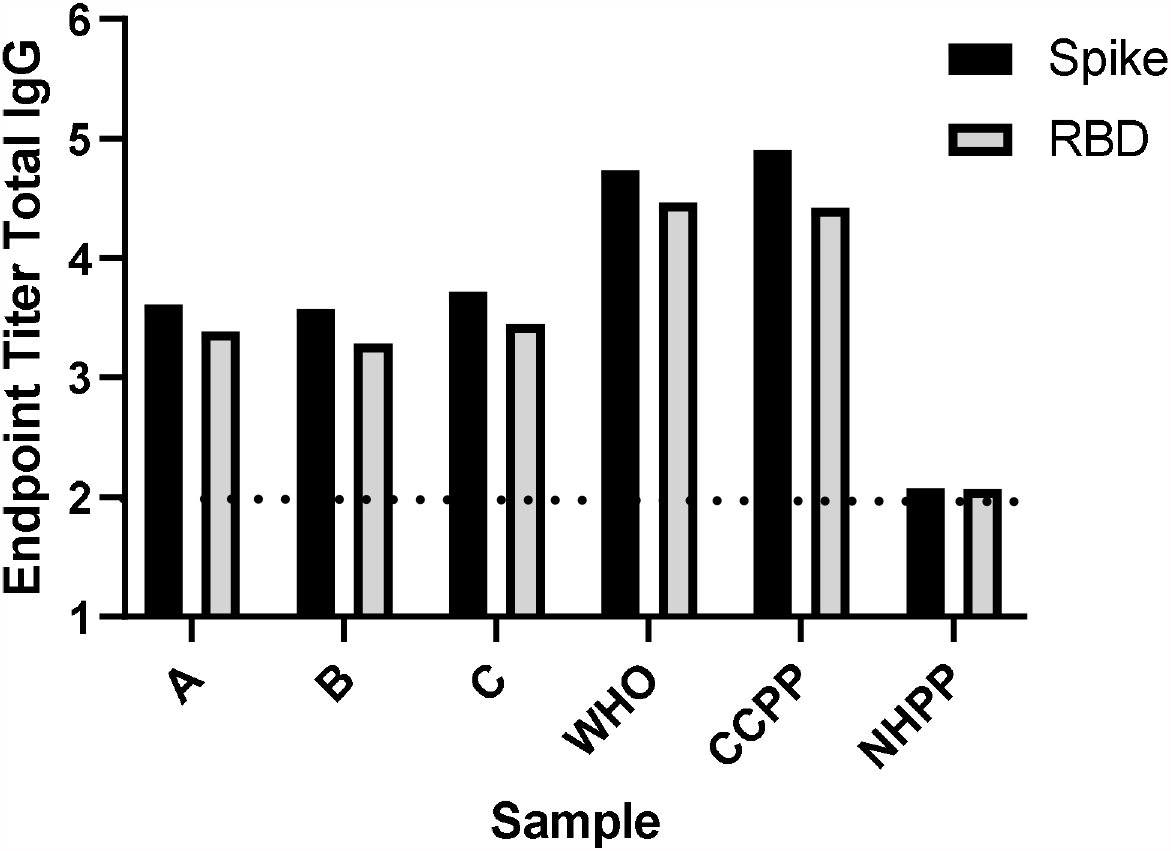
SARS-CoV-2 Spike and RBD Total IgG Endpoint Titers. evaluated for three individual samples (A, B, C) as well as positive WHO reference sample, pooled positive (CCPP) and negative (NHPP) control samples.

### 3.2 Pseudovirus neutralization

#### 3.2.1 Precision

Assay repeatability was assessed in terms of run to run variation. Samples from 2 subjects and 2 pooled plasma controls (CCPP and NHPP) were run by a single operator in duplicate across three separate days (*interday*), by a single operator in 3 different runs on the same day (*intraday*) and by three individual operators on the same day (*interoperator*). Data from individual samples (Positive, Negative) and pooled plasma controls (CCPP, NHPP) for all intraday, interday, and interoperator precision tests are shown in **Figure 7A, 7B**, and **7C**, respectively. Data from all 3 precision tests are compiled in **Figure 7D**. In each individual test and in the compiled data shown in **Figure 7D**, all samples fall within 2-fold of the geometric mean, demonstrating the assay is reproducible and precise.

**Figure 7:**
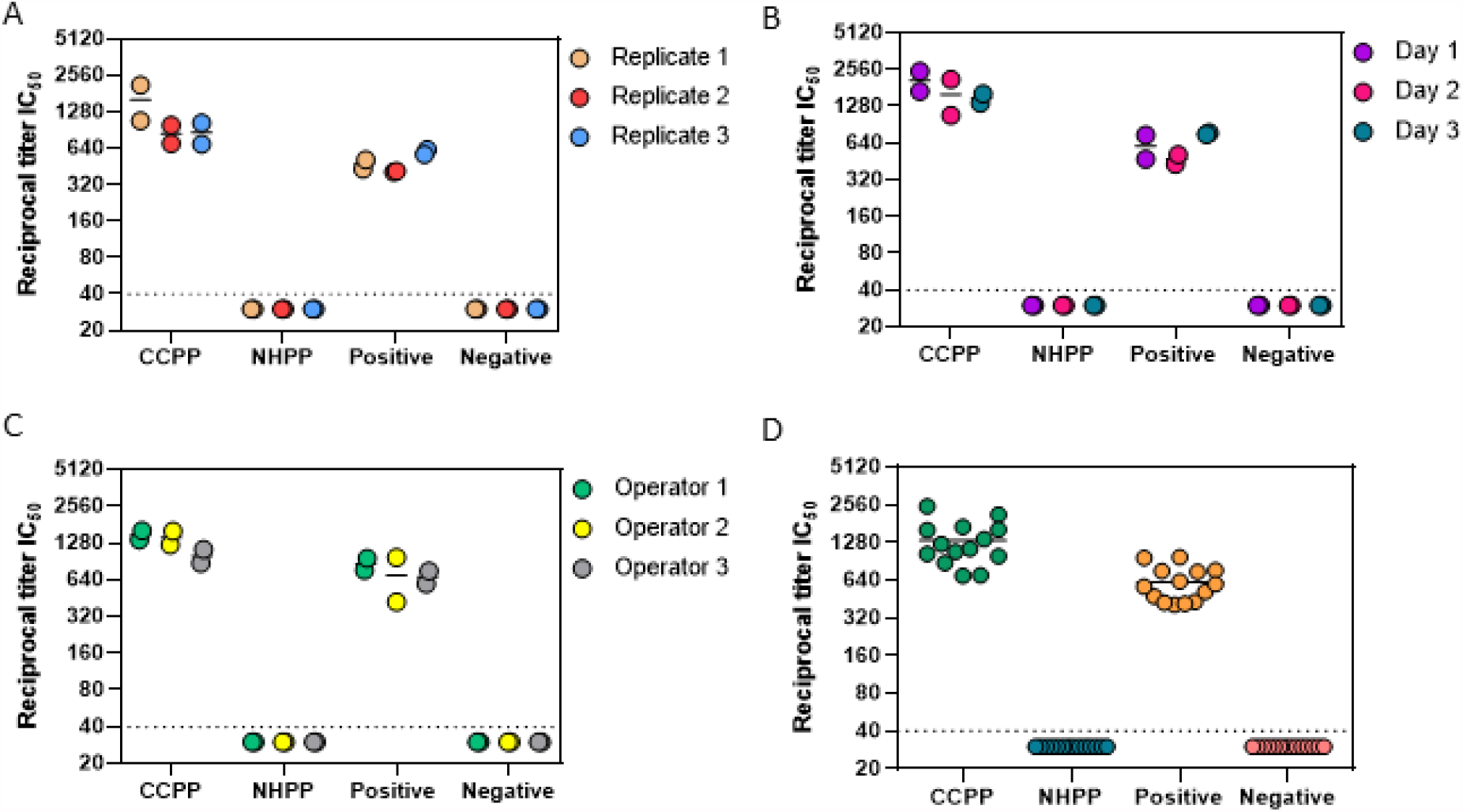
Pseudovirus Neutralization Reciprocal Titer IC_50_ Values. evaluated for two individual samples (Positive, Negative) as well as pooled plasma positive (CCPP) and pooled plasma negative (NHPP) control samples. Data shows intraday (A), interday (B), and interoperator (C) variability. Panel D represents IC_50_ values from all qualification runs combined.

#### 3.2.2 Linearity

RLU values from the Positive plasma samples of the interoperator precision run were used to evaluate assay linearity using linear regression analysis between Log-transformed RLU values and reciprocal plasma dilution concentrations. Only reciprocal plasma dilution concentrations within the linear range were used to generate representative graphs and calculate R^2^ values (**Figure 8**). As the plasma became more dilute, the PFU values plateaued, and the assay was no longer in the linear range (data not shown). Duplicate samples for each operator were averaged for an R^2^ of 0.939, 0.929, and 0.975 for Operators 1, 2, and 3, respectively. The adjusted multiple R^2^ value across all positive samples in all qualification runs came to 0.896, demonstrating the pseudovirus neutralization assay methodology is linear.

**Figure 8:**
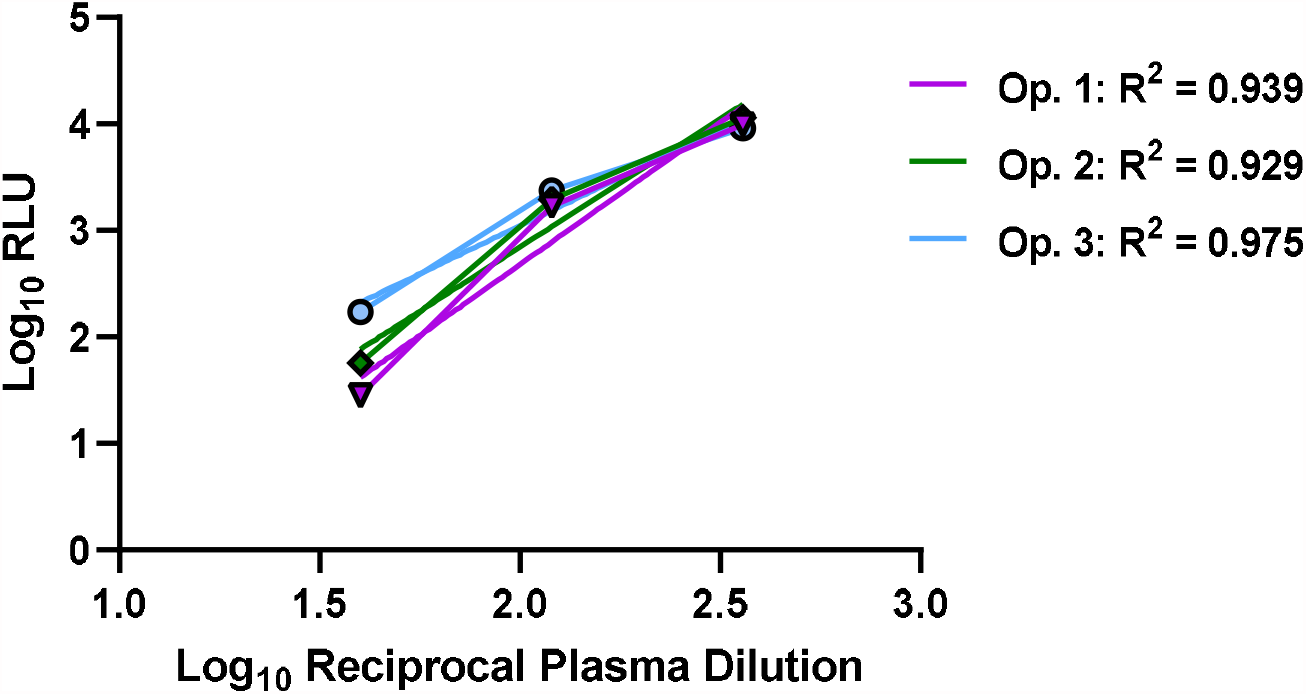
Graphical Representations of Linear Regression. models for the Positive control sample for each of the 3 operators. The calculated R^2^ value for each operator is shown in the legend.

#### 3.2.3 Specificity

Across our robust negative sample screen (pre-COVID-19 samples) for specificity, all 20 samples (100%) were negative for SARS-CoV-2 pseudovirus neutralization. All four positive samples (100%) inhibited SARS-CoV-2 pseudovirus by ≥ 50%. Neither of the two tested COVID-convalescent samples inhibited the VSV G pseudovirus by 50% at any plasma concentration (**Figure 9**), demonstrating the specificity of the assay for SARS-CoV-2 Spike antigen and not overlapping with an unrelated viral antigen.

**Figure 9:**
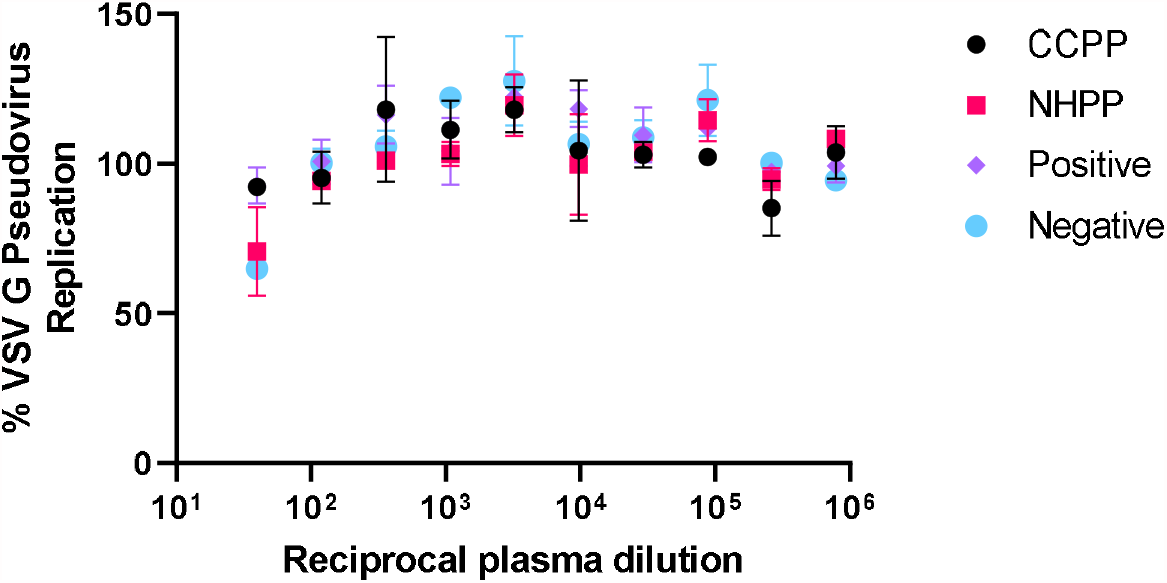
Inhibition of VSV G Pseudovirus Entry and Replication in HEK293T-ACE2 cells as a function of plasma concentration from healthy human donors (NHPP and Negative) or COVID-19 convalescent plasma samples (CCPP and Positive).

### 3.3 PRNT

#### 3.3.1 Precision

To examine the sources of variation present in the assay, we determined the geometric mean of the PRNT_80_ values for each sample within each category of precision evaluated (run, operator, day). Data from individual samples (Positive, Negative) and pooled plasma controls (CCPP, NHPP) for all intraday, interday, and interoperator precision tests are shown in **Figure 10A, 10B**, and **10C**, respectively. Data from all 3 precision tests are compiled in **Figure 10D**. In each individual test and in the compiled data shown in **Figure 10D**, all samples fall within 2-fold of the geometric mean, demonstrating the assay is reproducible and precise.

**Figure 10:**
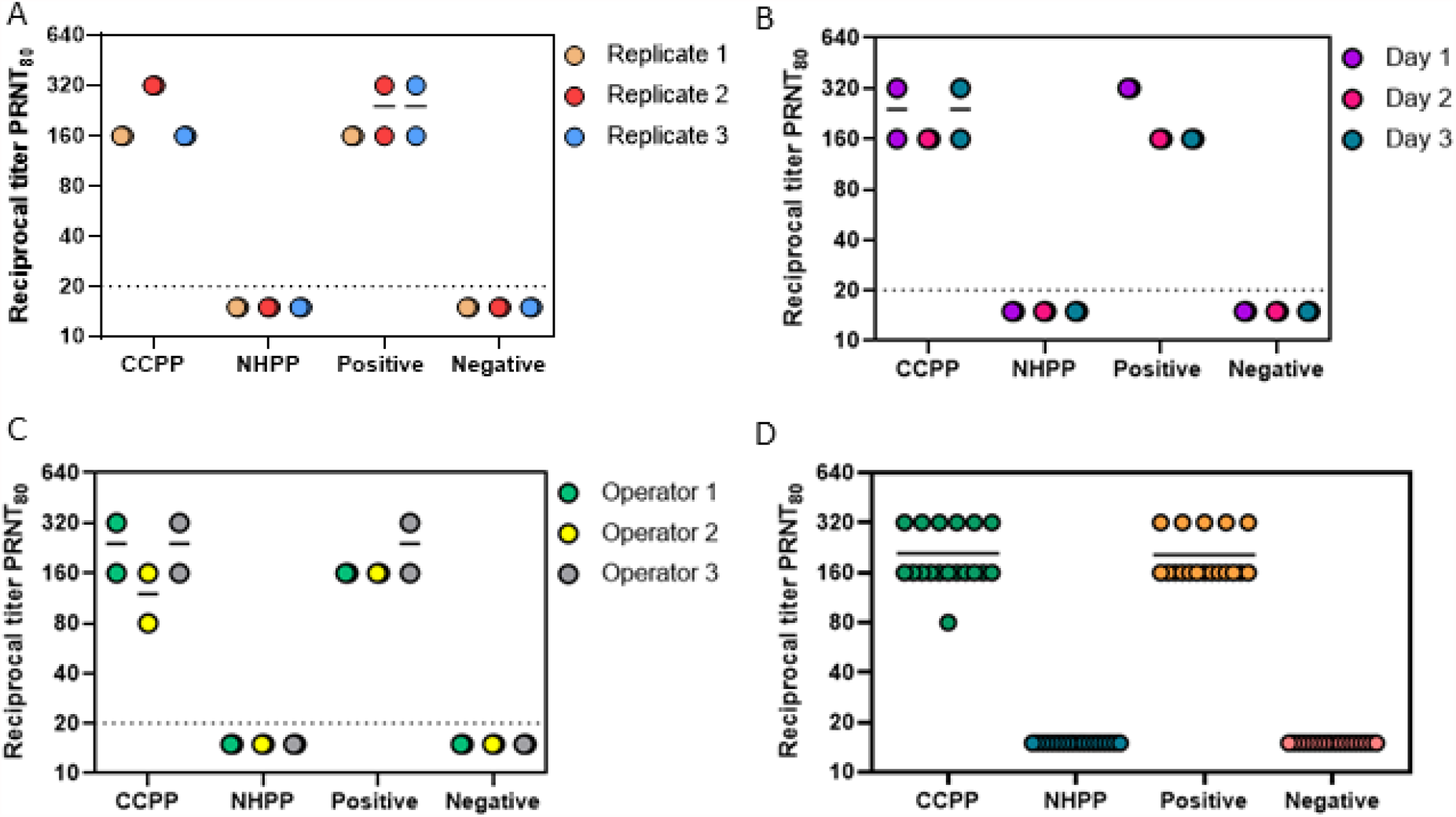
Reciprocal Titer PRNT_80_ Values. evaluated for two individual samples (Positive, Negative) as well as pooled plasma positive (CCPP) and negative pooled plasma (NHPP) control samples. Data show intraday (A), interday (B), and interoperator (C) variability. Panel D represents PRNT_80_ values from all qualification runs combined.

#### 3.3.2 Linearity

We evaluated the ability of the assay to return PFU values that were proportional to the dilution being examined using linear regression. Only the first four dilutions from each operator (1:80, 1:160, 1:320, and 1:640) were used to generate representative graphs and calculate R^2^ values (**Figure 11**). As the plasma became more dilute, the PFU values plateaued, and the assay was no longer in the linear range (data not shown). Positive control samples from each of the three operators were assessed using a simple linear regression. Each operator’s positive sample demonstrated a linear relationship (R^2^ = 0.922, 0.965, 0.950) between Log PFU and Log reciprocal dilution (**Figure 11**). These data demonstrate that the PRNT assay generates results that are linear with respect to sample dilutions and that this feature is consistent across assay operators.

**Figure 11:**
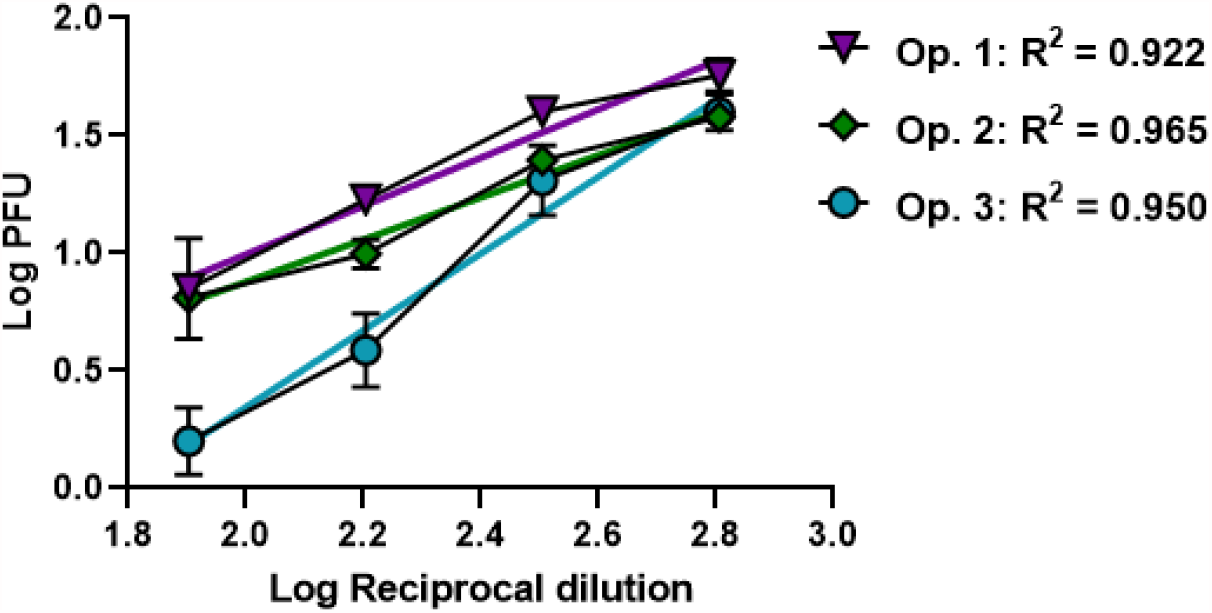
Graphical Representation of Linear Regression. models for the Positive control sample run by each of the 3 operators. The adjusted multiple R^2^ value across this sample set is 0.9157.

#### 3.3.3 Specificity

We next evaluated the specificity of serum/plasma samples to SARS-CoV-2 using a robust number of negative samples. Across our negative sample screen (pre-pandemic samples) for specificity, all 20 samples (100%) were negative for SARS-CoV-2 neutralization. All four positive samples (100%) inhibited SARS-CoV-2-PFUs by ≥ 80%. Both of these metrics meet our required endpoints for qualification of specificity of the methodology.

### 3.4 Assay Alignment

Many researchers have demonstrated that higher magnitude antibody responses often correlate with enhanced protection in endpoint assays^43-46^. Here we aimed to compare that between our own suite of qualified assays. Pooled plasma controls (CCPP and NHPP) were used across all assays and a single convalescent plasma sample was use in the pseudovirus neutralization and total IgG ELISA for RBD and Spike responses. We used these overlapping samples to demonstrate a correlation between EPT values and neutralization for High, Medium and Low responder samples (**Figure 12, Supplementary Figure 5**). We observed that CCPP consistently returned High responder values, the single convalescent samples consistently returned Medium responder values and NHPP was consistently Low across all endpoint assays evaluated (**Supplemental Figure 5**). Furthermore, we are able to demonstrate a significant correlation between Reciprocal Titer IC_50_ from pseudovirus neutralization and both Total IgG Spike EPT (adjusted R^2^ = 0.9623) and Total IgG RBD EPT (adjusted R^2^ = 0.8921), as well as a correlation between Total IgG for Spike and RBD EPT (adjusted R^2^ = 0.9814) (**Figure 12**).

**Figure 12:**
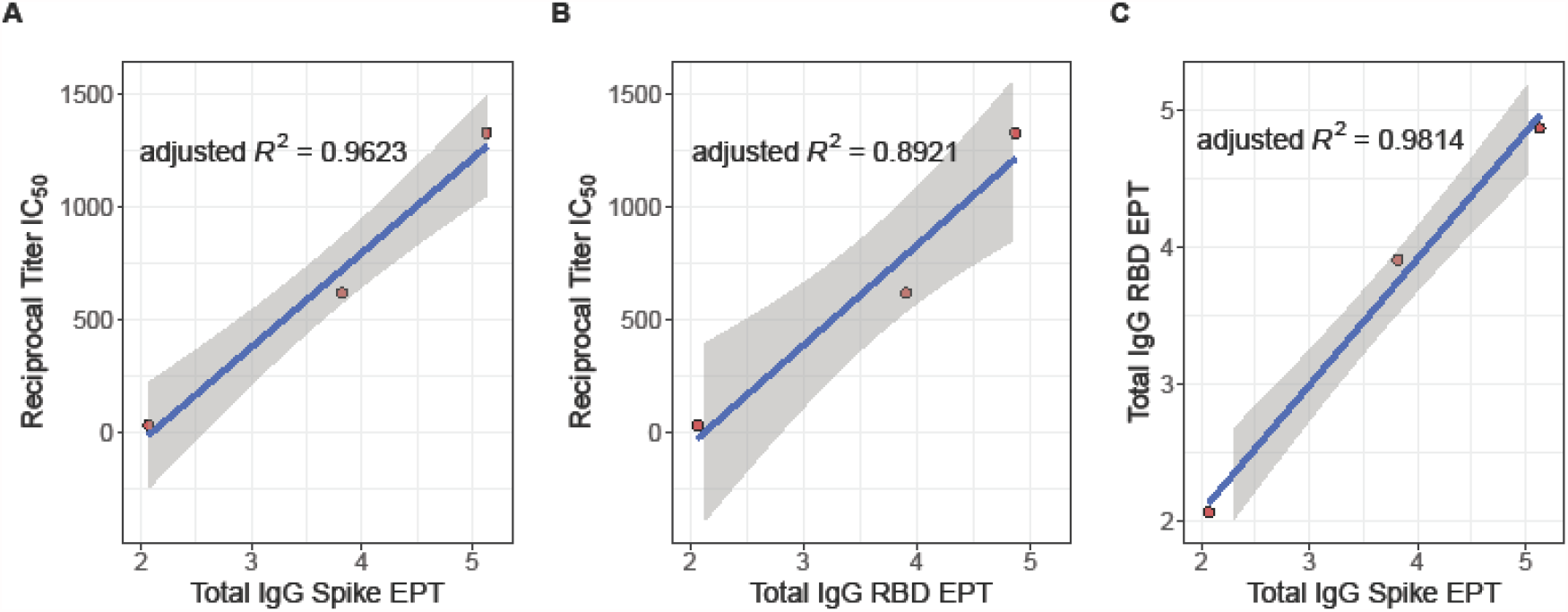
Total IgG EPT Correlates with Pseudovirus Neutralization IC_50_ values. Correlations between Reciprocal Titer IC_50_ and (A) Total IgG Spike EPT, (B) Total IgG RBD EPT and (C) Total IgG RBD EPT and Total IgG Spike EPT. In each panel the blue line represents the line of best fit and shaded grey is the 95% confidence interval. Adjusted R^2^ values shown for each correlation.

## 4 Conclusions

Here we have described robust methodologies qualified to evaluate the essential protective features of the humoral response against SARS-CoV-2. The EPT ELISA, pseudovirus neutralization and PRNT assays were all determined to be precise, linear, and specific for antibody-specific SARS-CoV-2 responses. These assays can be reliably used across days and by different trained operators to measure the effectiveness of clinical vaccine candidates both in the pipeline and approved for emergency use. In addition, the use of a WHO reference standard makes these qualified assays more accessible to other labs and researchers for use in comparing SARS-CoV-2 vaccine candidates.

## Acknowledgements

The authors would like to express their gratitude to Seattle Children’s Research Institute Leadership for their guidance, compassion, and mentorship. The authors would like to thank the Harrington Lab at Seattle Children’s Research Institute as well as our colleague participants in the Center for Global Infectious Disease Research Biorepository for their commitment to science and resilience during a pandemic. The authors would like to thank their long-time colleague Dr. Jesse Erasmus for invaluable technical help when adopting and optimizing neutralization assays.

The datasets generated and/or analyzed during the study presented here are available from the corresponding authors on reasonable request.

Research reported here was supported by: National Institute of Allergy and Infectious Diseases (NIAID) of the National Institutes of Health under award numbers 1UM1AI148373-01 and 3UM1AI148373-01S1, a Bill & Melinda Gates Foundation (BMGF) grant to N.P.K. (OPP1156262), and B.P.B is supported through award number F32HD102290 from the National Institute of Child Health and Human Development (NICHD). The content is solely the responsibility of the authors and does not necessarily represent the official views of the National Institutes of Health or BMGF.

## Author Contribution Statement

SEL, BJB, TP, EC and RNC designed experiments; SEL, BJB, TP, EC, BW, and EJ carried out experiments and SEL, PQ, BJB and BPB executed statistical analysis; LC, SW, EK, CS and NPK provided reagents; SEL, BJB and EC wrote the manuscript and prepared figures and tables with assistance from PQ, BPB, SLB, and RNC. All authors reviewed the manuscript.

## Additional Information

All authors declare no financial or alternative competing interests with the data described in this manuscript.

