## Supplemental Figures for "Qualification of ELISA and neutralization methodologies to measure SARS-CoV-2 humoral immunity using human clinical samples"

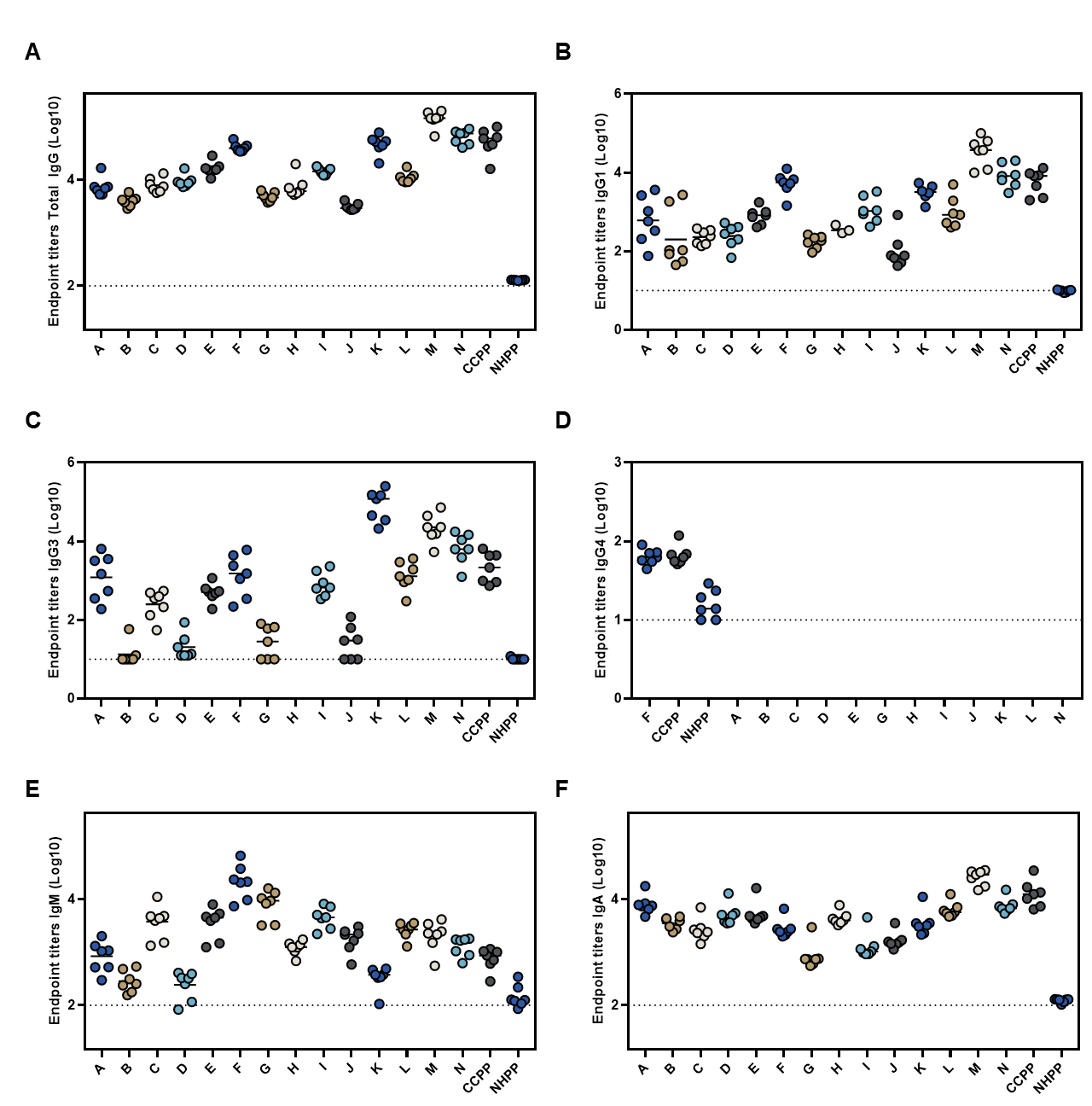

**Supplemental Figure 1: Cumulative EPT across samples from ELISA Precision analysis.** Log 10 cumulative EPT values across individual samples and controls (CCPP, NHPP) for SARS-CoV-2 spike-antigen specific antibody responses by class. A) Total IgG, B) IgG1, C) IgG3, D) IgG4, E) IgM, and F) IgA.

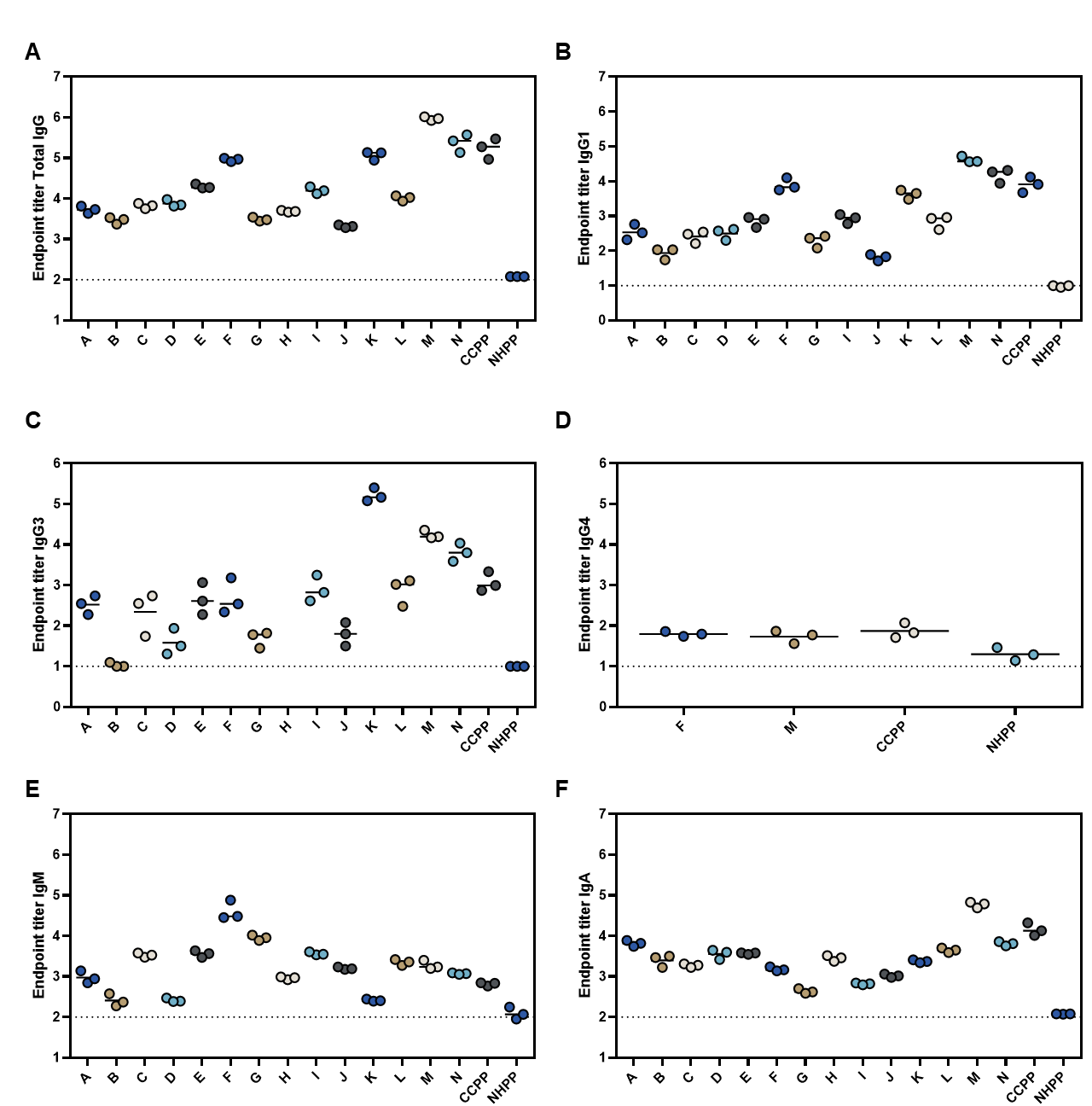

**Supplemental Figure 2: Intraday EPT across samples from ELISA Precision analysis.** Log 10 EPT values from Intraday analysis across individual samples and controls (CCPP, NHPP) for SARS-CoV-2 spike-antigen specific antibody responses by class. A) Total IgG, B) IgG1, C) IgG3, D) IgG4, E) IgM, and F) IgA.

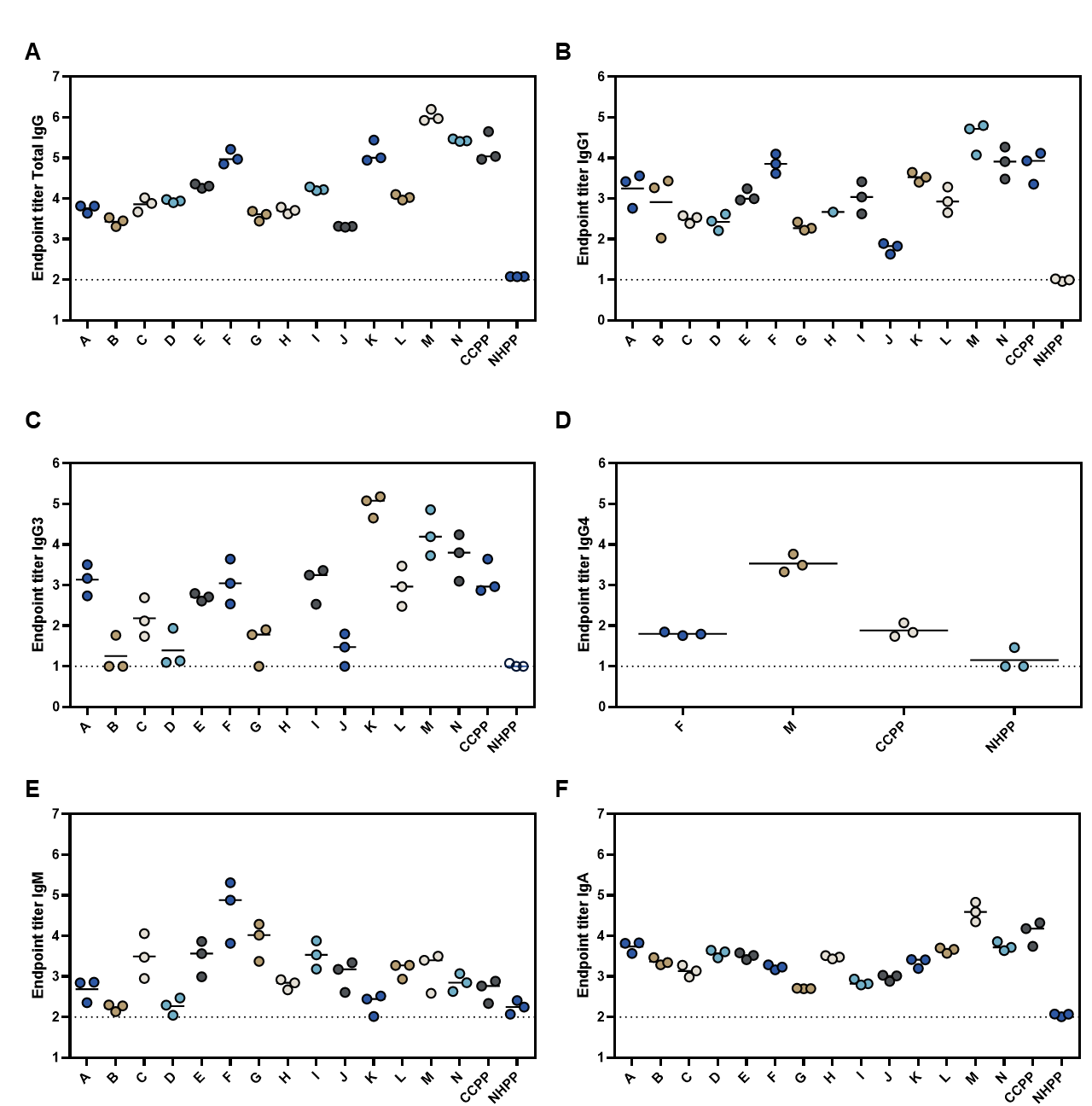

**Supplemental Figure 3: Interday EPT across samples from ELISA Precision analysis.** Log 10 EPT values from Interday analysis across individual samples and controls (CCPP, NHPP) for SARS-CoV-2 spike-antigen specific antibody responses by class. A) Total IgG, B) IgG1, C) IgG3, D) IgG4, E) IgM, and F) IgA.

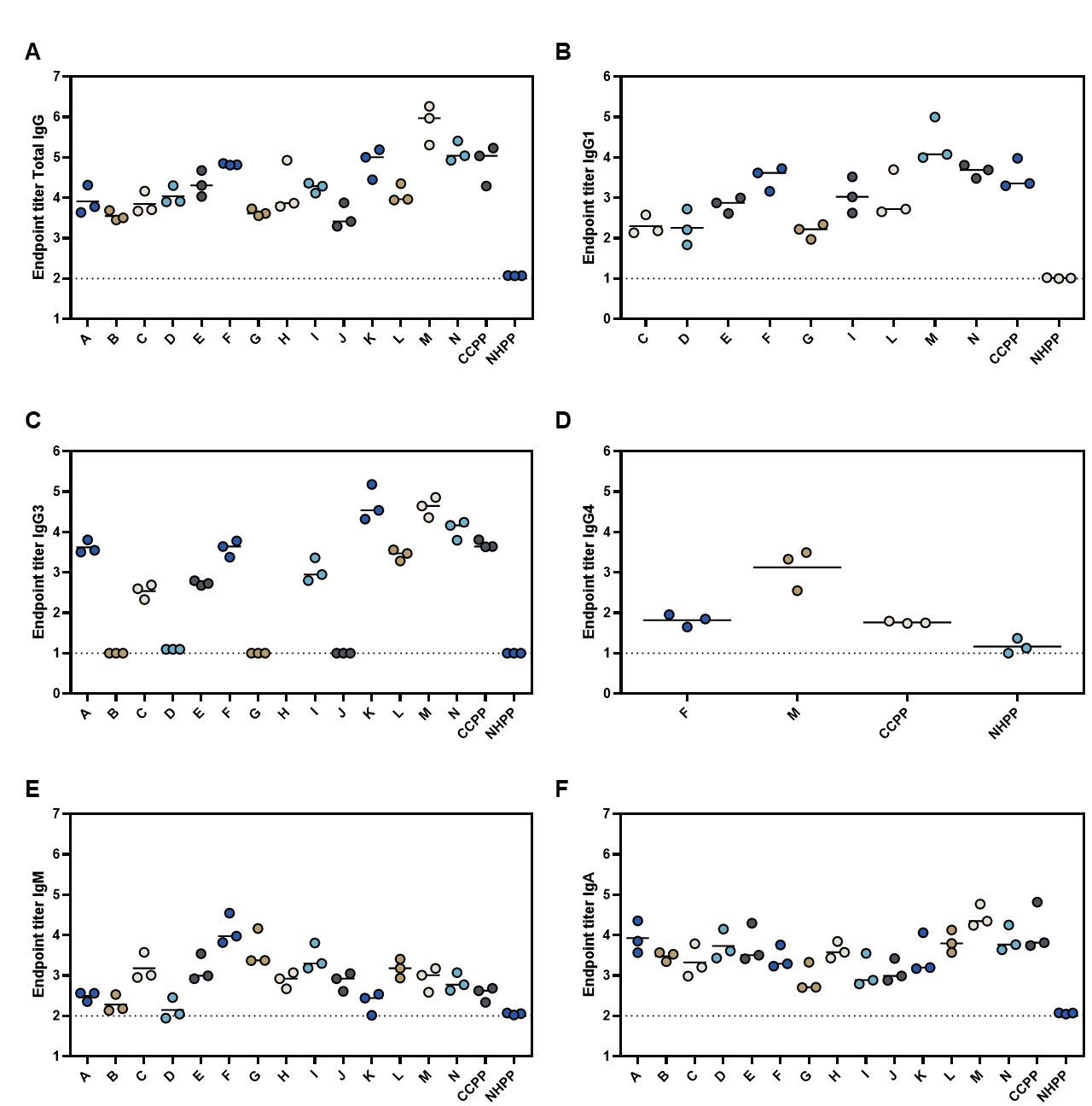

**Supplemental Figure 2: Interoperator EPT across samples from ELISA Precision analysis.** Log 10 EPT values from Interoperator analysis across individual samples and controls (CCPP, NHPP) for SARS-CoV-2 spike-antigen specific antibody responses by class. A) Total IgG, B) IgG1, C) IgG3, D) IgG4, E) IgM, and F) IgA.
